# Inadequate antibody validation places substantial numbers of animal and human tissue samples at risk of waste

**DOI:** 10.64898/2026.02.19.706766

**Authors:** Michael Biddle, Jemma Cooper, Katherine Blades, Dominic Ruddy, Eva M. Krockow, Harvinder Virk

## Abstract

Lack of antibody validation by researchers frequently misdirects biomedical research, yet the ethical consequences — particularly avoidable use of animal and human biological materials — remain unquantified. Using focus groups (n=12), surveys (n=107), and systematic analysis of 785 publications, we examined how researchers select and validate antibodies, and quantified the downstream ethical costs. Antibody selection is substantially influenced by social factors such as previous laboratory use and peer recommendations, while systematic evaluation of performance characteristics remains limited. From a dataset of 614 antibodies subjected to rigorous characterisation using knockout controls, 97 (15.8%) failed across all tested applications; we systematically searched for publications linked to these antibodies. Within the 760 publications (those where validation status could be confirmed), only 120 (15.8%) presented any validation evidence — despite 72.0% of surveyed researchers reporting having used at least one recommended validation method. The remaining 640 papers reported a minimum of 8,064 animal samples and 4,424 human tissue samples used with antibodies of demonstrated poor performance and without evidence of fitness for the specific experimental purpose; we describe these as samples at risk of waste. Reserving the term waste for samples used with reagents since withdrawn from sale, and therefore irreproducible, conservative extrapolation gives lower-bound, order-of-magnitude estimates of millions of animal and human tissue samples consumed globally. Researchers identified time, cost, and lack of supervisor support as primary barriers, whilst strongly supporting open data sharing, dedicated validation funding, and publisher requirements as solutions. These findings provide a first quantification of the ethical costs of inadequate antibody validation and highlight the need for coordinated stakeholder interventions to reduce avoidable biological sample waste in biomedical research.

## INTRODUCTION

Antibodies are critical reagents that enable researchers to detect, quantify, and isolate specific proteins within complex biological samples. However, research antibodies do not always bind their intended targets, or may bind additional unintended targets [1]. Poor antibody specificity has repeatedly misdirected biomedical research across diverse fields, leading to incorrect conclusions about protein function and localisation, unreliable biomarker discovery, and misidentification of drug targets [2–10]. These failures have been documented across multiple disciplines, highlighting a persistent and systemic issue rather than isolated occurrences.

Evidence suggests that problems with antibody data reporting and quality may represent a significant contribution to the expensive and frequent replication failures seen in preclinical research [11]. The estimated economic waste is in the region of 1 billion USD per annum in the US alone, excluding the substantial opportunity cost of misdirected research effort [12,13]. The underlying reasons for this research system failure are complex, driven both by difficulties in producing and characterising high-quality antibodies and by gaps in researcher awareness, education, and incentives [14,15].

To date, researchers attempting to quantify the scale of this problem have focused on the impact on literature reliability and economic waste [1,11]. It is clear, however, that there is also an ethical dimension. This includes not only the waste of animals in antibody production [16], but also the waste of biological samples—both animal-derived and donated by humans—in research that relies on antibodies not fit for the specific purpose. Despite increasing attention to the reproducibility and economic impact of poor antibody performance, the ethical consequences remain largely unquantified.

Previously, we systematically assessed the selectivity of antibodies within three commonly used research applications across a sample of over 600 antibodies directed at neuroscience-relevant proteins [1]. We found a high prevalence of poorly performing antibodies under the standardised, industry-endorsed protocols used [17]. We also observed that, at least for immunocytochemistry–immunofluorescence (ICC-IF), presentation of antibody validation data within the published literature was very limited [1]. That analysis suggested that uptake of best-practice recommendations from the International Working Group for Antibody Validation (IWGAV) [18] in the published literature is very limited: relevant antibody validation data was not presented in 98 of 112 publications utilising antibodies that had performed poorly in ICC-IF.

Against this backdrop, there is a clear need to understand how researchers choose and validate antibodies in practice, and to quantify the downstream ethical cost of suboptimal decisions. We designed a mixed-methods approach combining qualitative focus groups and a semi-quantitative survey of researchers to understand their decisions related to antibody selection and validation. We subsequently contrasted the self-reported results with observed behaviour as evidenced by reporting of antibody choice and use in publications. We adopted this between-method triangulation to cross-check evidence generated from different sources and methodological approaches.

## RESULTS

### Focus group participants

We recruited an opportunity sample of 12 UK-based researchers or research students with experience of using antibodies. The sample consisted of 5 males, 6 females, and 1 identifying as ‘Other’, with a mean age of 26.7 years (SD = 5.5 years). Eight participants were PhD students, with additional representation from one undergraduate researcher, a technician with extensive laboratory experience, one participant working in clinical trials, and one postdoctoral researcher. Participants reported experience with a range of antibody-based methods, including Western blotting, immunohistochemistry, and flow cytometry.

### Survey participants

A total of 107 researchers and research students with confirmed institutional emails and experience using antibodies completed the online survey. Respondents represented a range of experience levels: 7 (6.5%) at undergraduate level, 2 (1.9%) at Masters level, 43 (40.2%) PhD students, 15 (14.0%) postdoctoral or early career researchers, 13 (12.1%) experienced postdoctoral researchers, 23 (21.5%) independent investigators, and 4 (3.7%) research staff (e.g., research assistants/scientists).

Laboratory experience ranged widely: 7 (6.5%) reported less than 1 year, 37 (34.6%) between 1– 5 years, 21 (19.6%) between 5–10 years, and 42 (39.3%) more than 10 years. The majority of respondents (92/107; 86.0%) reported having direct experience selecting antibodies for research purposes. They also reported using a variety of antibody-based methods, including western blotting (84), immunohistochemistry (60), flow cytometry (57), ELISA (52), immunocytochemistry (45), and immunoprecipitation (39), with smaller numbers using methods such as protein arrays, chromatin immunoprecipitation, and immunoelectron microscopy. The cohort was concentrated in the UK (81 of 107; 76%), with further respondents from Canada (9), the United States (8) and China (4), single respondents from Belgium, France and Taiwan, and two from an industrial setting (full survey responses are available in our Zenodo deposit). Because the demographics of the global antibody-using community are unknown, we make no claim of statistical representativeness and present the survey as exploratory and corroborative rather than prevalence-estimating.

### How do researchers choose antibodies – qualitative themes

Three broad themes of factors influencing antibody selection emerged from focus group discussions: (1) social and reputational factors, (2) convenience and pragmatic concerns, and (3) engagement with validation data (Figure 1A). Illustrative quotes mapped to each theme and subtheme are provided in S1 Table.

**Figure 1.**
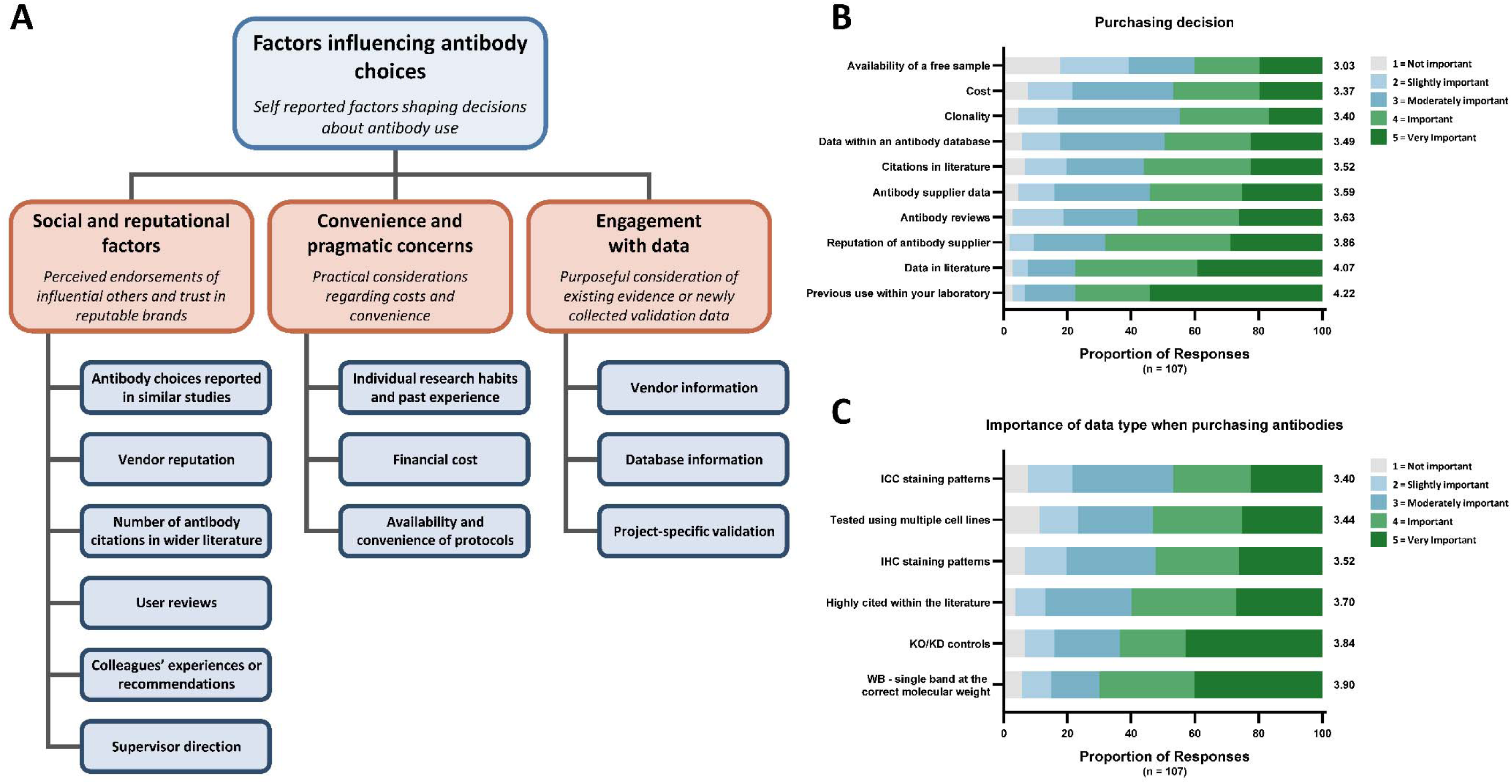
Factors influencing antibody selection among researchers. **(A)** Thematic map from focus group discussions (n=12 participants) identifying three categories of factors influencing antibody selection: social and reputational factors, convenience and pragmatic concerns, and engagement with data. **(B)** Survey responses (n=107) rating importance of factors in purchasing decisions on a 5-point Likert scale ordered by combined percentage rating as important or very important (ratings 4–5). Previous use within the laboratory (77.6%, 83 of 107 rating as important or very important) and data in literature (77.6%, 83 of 107) were rated most highly, followed by supplier reputation (68.2%, 73 of 107) and citations in literature (56.1%, 60 of 107). Mean Likert ratings are presented to the right of the bars. **(C)** Survey responses rating importance of validation data types in purchasing decisions. Western blot data showing a single band at correct molecular weight was rated most highly (70.1%, 75 of 107), followed by knockout/knockdown (KO/KD) controls (63.6%, 68 of 107) and highly cited antibodies (59.8%, 64 of 107). Mean Likert ratings are presented to the right of the bars. The data underlying this figure can be found in https://doi.org/10.5281/zenodo.18682587.

Participants consistently described using social proof and reputation as guides for antibody selection, particularly consulting published literature to identify which antibodies other researchers had used, with citation counts serving as a primary selection criterion. Colleagues and supervisors strongly influenced choices, with junior researchers often following supervisor direction with minimal independent evaluation: “I’ve just been told what to pick. […] I’ve never even heard of the word validating antibodies until […] this meeting” (Participant 12; S1 Table).

Established habits and past experience shaped decisions, with many researchers choosing antibodies simply because they had previously used them. Financial considerations influenced choices in complex ways—while most sought affordable options, one participant explicitly used price as a quality indicator (S1 Table).

Participants reported consulting validation data from vendor websites and independent databases, evaluating features such as Western blot band clarity as indicators of quality (S1 Table).

### How do researchers choose antibodies – quantitative survey responses

Survey responses quantified the relative importance of these factors. For each factor assessed, ratings spanned the full range from not important to very important, indicating substantial diversity in what individual researchers prioritize when selecting antibodies (Figure 1B–C).

Previous use within the laboratory and data in literature were rated as the most important factors, with 77.6% of respondents rating each as important or very important (scores 4–5 on a 5-point scale), though previous use received more ‘very important’ ratings. Reputation of antibody supplier was rated as important or very important by 68.2% of respondents, and citations in literature by 56.1%. Cost was rated as important or very important by 46.7%, while availability of a free sample was rated as important or very important by 40.2%. Clonality (whether monoclonal or polyclonal) received the lowest importance rating at 44.9% (Figure 1B).

When asked specifically about different types of validation data, survey respondents rated Western blot data showing a single band at the correct molecular weight as most important (70.1% rating as important or very important), followed by knockout/knockdown controls (63.6%), and high citation counts within the literature (59.8%). Staining pattern data for immunohistochemistry (52.3%) and immunocytochemistry (46.7%), and testing using multiple cell lines (53.3%) received relatively lower ratings (Figure 1C).

### How do researchers validate antibodies – quantitative survey responses

Only 36 of 107 survey participants (33.6%) reported awareness of the IWGAV [18] five-pillar framework of antibody validation. We categorised participants’ self-reported validation methods according to this framework. Participants reporting use of genetic strategies (CRISPR knockout or siRNA/shRNA knockdown), tagged expression validation, independent antibodies, immunocapture mass spectrometry, or orthogonal validation including use of both positive and negative control cell lines were classified as using IWGAV-aligned validation approaches [18]. We adopted the looser operational definition of orthogonal validation used by many antibody vendors, under which comparison of antibody signal across positive and negative control cell lines is treated as orthogonal; under a strict, antibody-independent definition (counting only transcriptomic or comparable orthogonal assays), the proportion of respondents reporting at least one IWGAV-aligned method falls from 72.0% to 58.9%, although the conclusion that most respondents rely on a single, often basic, approach is unchanged. Responses merely indicating technical controls (fluorescence minus one, isotype controls, secondary-only controls, blocking peptides, or single control lines) without IWGAV-aligned methods were coded as technical controls rather than validation.

Overall, 77 of 107 participants (72.0%) reported using at least one IWGAV-aligned validation method, while 30 (28.0%) reported no IWGAV-aligned validation—of whom 26 used only technical controls and 4 reported no validation methods of any kind (Figure 2A). Among the 77 participants using IWGAV-aligned validation, 44 (57.1%) reported using a single pillar, 22 (28.6%) used two pillars, 10 (13.0%) used three pillars, and 1 (1.3%) reported using five pillars. Among the 26 participants using only technical controls, 21 (80.8%) reported using a single control type.

**Figure 2.**
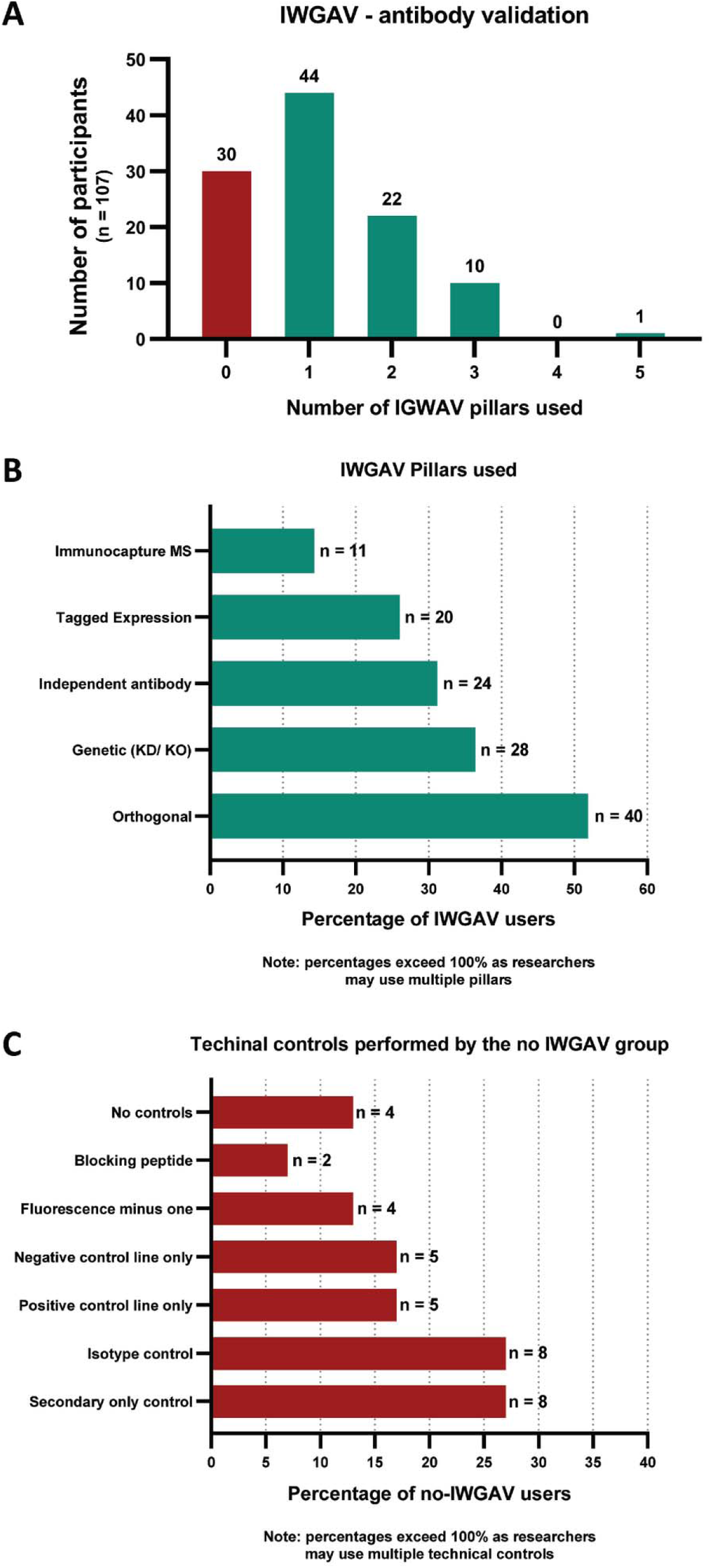
Self-reported antibody validation practices among researchers. (A) Distribution of IWGAV pillar usage among survey respondents (n=107). Of 107 respondents, 77 (72.0%) used at least one IWGAV-aligned validation strategy, while 30 (28.0%) used no IWGAV pillars, relying instead on technical controls only (n=26) or performing no validation (n=4). Among IWGAV users, 44 used a single pillar, 22 used two pillars, 10 used three pillars, and 1 used five pillars. (B) Specific IWGAV pillars used among 77 IWGAV-aligned researchers. Orthogonal validation was most common (n=40, 51.9%), followed by genetic strategies (n=28, 36.4%), independent antibodies (n=24, 31.2%), tagged expression (n=20, 26.0%), and immunocapture MS (n=11, 14.3%). Researchers using both positive and negative control cell lines were classified as orthogonal validation. Percentages exceed 100% as researchers may use multiple pillars. (C) Technical controls used among the 30 respondents not using IWGAV-aligned validation, including 4 who performed no controls of any kind. Secondary-only controls and isotype controls were each most common (n=8), followed by positive or negative control cell lines used alone (n=5 each), fluorescence minus one (n=4), and blocking peptide (n=2). The data underlying this figure can be found in https://doi.org/10.5281/zenodo.18682587.

Orthogonal validation was the most commonly reported IWGAV-aligned approach (40 participants, 51.9%), followed by genetic strategies (28 participants, 36.4%), independent antibodies (24 participants, 31.2%), tagged expression (20 participants, 26.0%), and immunocapture mass spectrometry (11 participants, 14.3%) (Figure 2B). Among the 30 respondents not using IWGAV-aligned validation, secondary-only controls and isotype controls were most prevalent (each 8 participants, 26.7%), followed by positive control line only and negative control line only (each 5 participants, 16.7%), fluorescence minus one (4 participants, 13.3%), and blocking peptide (2 participants, 6.7%); 4 participants (13.3%) reported no controls of any kind (Figure 2C).

### How do researchers validate antibodies – observations from published literature

To understand how antibody validation is practised in published research, we examined the literature associated with antibodies that had failed independent characterisation. The YCharOS consortium tested 614 commercial antibodies against their target proteins using standardised, knockout-controlled protocols across western blot, immunoprecipitation and immunofluorescence; 97 (15.8%) failed in every application tested [1,17]. From these 97 antibodies, literature searches identified publications for 35 (see Methods), and we analysed 785 papers associated with them. Of these, 19 were excluded as sample numbers could not be extracted, leaving 766 papers for biological sample analysis. Of the 35 antibodies, 16 had been discontinued by their manufacturers at the time of analysis. Details of antibody selection and literature search methods are provided in the Methods section.

Of the 766 papers with extractable sample data, validation status could be determined in 760; the remaining 6 contained usable sample data but could not be classified by validation status and are included in sample totals only. Across the 760 papers where validation status could be determined, only 120 (15.8%) presented some form of antibody validation data aligned with the IWGAV five pillars (Figure 3A); the remaining 640 papers (84.2%) reported no validation evidence. Among the minority of papers presenting validation (n=120), knockdown was the most commonly reported approach (83 papers, 69.2%), with the definitive knockout control used in only 20 papers (16.7%), followed by overexpression or tagged protein validation (40 papers, 33.3%). Only 1 paper (0.8%) used independent antibodies for validation (S3 Table). Individual papers could present multiple validation approaches.

**Figure 3.**
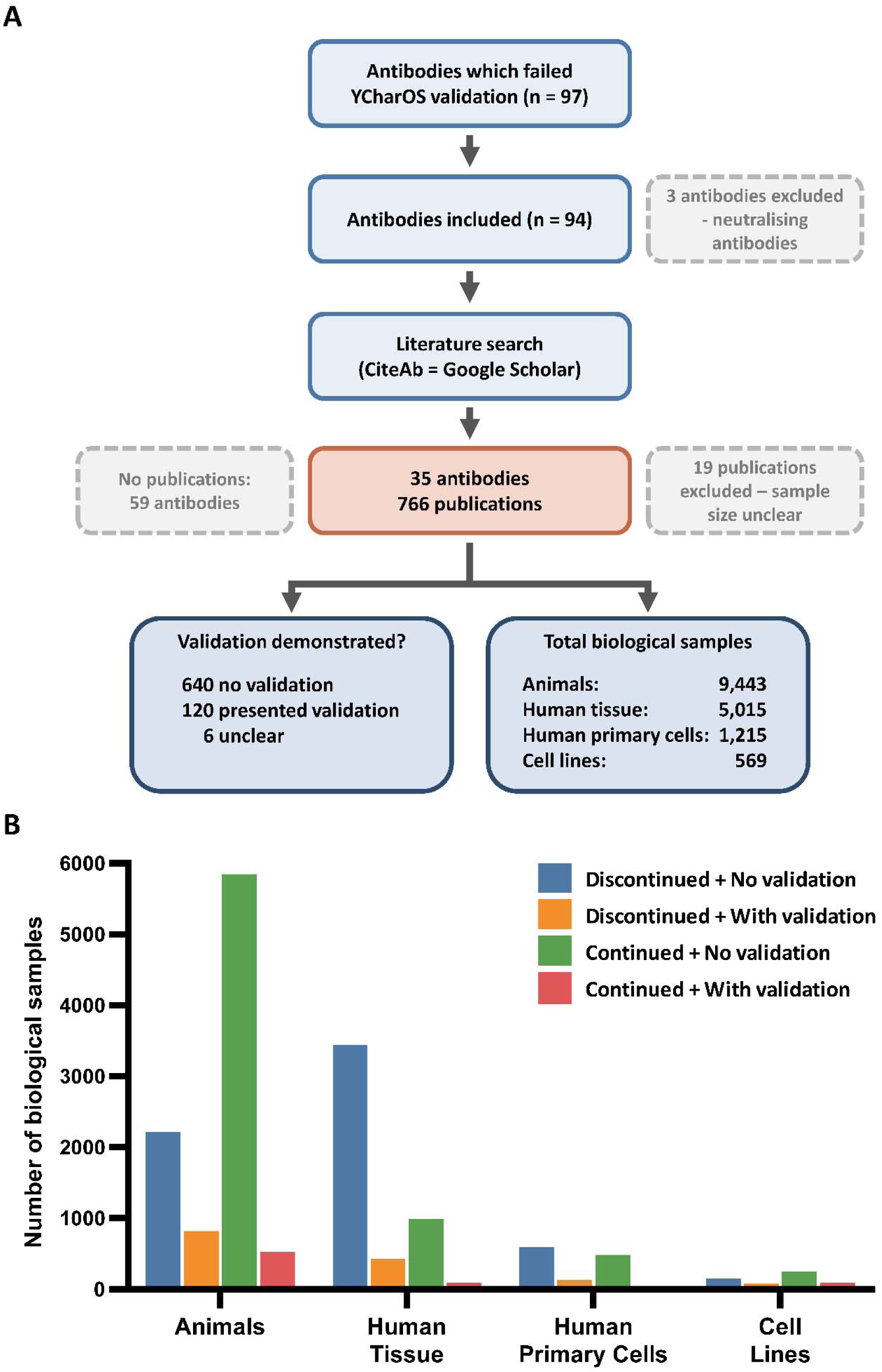
Literature analysis workflow and biological sample waste associated with insufficient antibody validation. (A) Flowchart showing selection of publications for analysis. From 97 antibodies failing YCharOS validation testing, 3 neutralising antibodies were excluded. Literature searches identified publications for 35 antibodies (16 discontinued at time of analysis); 59 had no associated publications. A total of 785 publications were reviewed; 19 were excluded as sample numbers could not be extracted, leaving 766 papers for biological sample analysis. Validation status could be determined in 760 of these papers (640 without validation, 84.2%; 120 with validation, 15.8%); the remaining 6 contained extractable sample data but could not be classified by validation status and are included in sample totals only. (B) Biological sample usage stratified by antibody commercial status and validation evidence. Sample totals in (A) exceed the sum of validated and non-validated categories as the 6 papers with unclear validation status contributed to overall sample counts but could not be allocated to either category. The data underlying this figure can be found in https://doi.org/10.5281/zenodo.18682587.

These approaches are not equivalent in stringency. Knockdown reduces the amount of target protein but does not remove it, so residual signal is difficult to interpret and a partial reduction is consistent with both specific and non-specific binding. Overexpression and tagged-protein approaches show that an antibody can detect the target when it is abundant, but not that it is selective for the target at the levels normally found in a cell. Knockout is the more definitive test, and it is the test these antibodies failed.

### What is the extent of biological sample use – observations from published literature

To quantify the ethical cost of insufficient antibody validation, we extracted sample numbers from publications where these were explicitly reported. Our literature analysis was restricted to publications available online from December 2000 to October 2025; earlier publications that may have used these antibodies were not captured.

Across 766 papers with extractable sample data, we identified a minimum of 9,443 animal samples (primarily mouse and rat, with additional samples from other species), 5,015 human tissue samples, 1,215 human primary cell samples, and 569 cell lines (Figure 3A, B; S2 Table). We included only explicitly reported sample numbers and excluded any studies where counts could not be determined from the methods or figure legends. Full sample counts are provided in S2 Table.

The overwhelming majority of biological sample consumption occurred in studies presenting no context-specific validation data. Across 640 papers without validation data, 8,064 animal samples, 4,424 human tissue samples, 1,081 human primary cell samples, and 400 cell lines were consumed (Figure 3B). The 120 papers presenting validation data used 1,339 animal samples, 513 human tissue samples, 134 human primary cells, and 168 cell lines.

Because papers may perform validation without reporting it, we tested how sensitive these totals are to silent validation. If 10%, 20% or 30% of the 640 papers without reported validation had in fact validated without reporting, the animal and human tissue samples at risk of waste would fall from 8,064 and 4,424 to 7,258 and 3,982, to 6,451 and 3,539, and to 5,645 and 3,097 respectively, so the global estimates remain in the millions at every rate. The waste estimate itself is unaffected by under-reporting, because it is constructed only from reagents that have since been withdrawn from sale, which renders the associated experiments irreproducible whether or not validation was performed.

Within the papers without validation data, the distribution of sample use differed by sample type and antibody commercial status. For human tissue, 3,437 of 4,424 samples (77.7%) in papers without validation data used discontinued antibodies, compared with 987 (22.3%) using commercially available antibodies. For animal samples, the pattern was reversed: commercially available antibodies accounted for 5,847 of 8,064 animal samples (72.5%), while discontinued antibodies accounted for 2,217 (27.5%). This difference likely reflects the higher number of publications using commercially available antibodies (396 papers vs. 244) and the concentration of discontinued antibodies in study areas using human tissue such as immunohistochemistry. Notably, all antibodies in this analysis had failed YCharOS characterization performed in human cell lines using knockout controls, meaning the high proportion of human tissue samples consumed with discontinued antibodies represents consumption with antibodies that demonstrably failed against human antigens.

Of the 35 antibodies in our analysis, 16 (45.7%) had been discontinued by their manufacturers and 17 (48.6%) were not recommended for use in mouse, with 6 overlapping both categories. In total, 27 of 35 antibodies (77.1%) carried at least one of these vendor-level flags, which are independent of the YCharOS standardised testing results. When weighted by sample use, the mouse-not-recommended antibodies account for only 8.5% (804 of 9,443) of the animal samples in our dataset, so this flag does not materially change the animal-sample estimate. The animal-sample inference instead rests on the demonstrated specificity failure and the absence of reported validation, and for the discontinued antibodies on their withdrawal from the market, which renders the associated experiments irreproducible regardless of species.

We extrapolated from our findings to estimate the global scale of biological sample waste (see Methods for full calculation). Applying the observed failure rate from our antibody panel (97/614 = 15.8%) to conservative global market size estimates (1–2 million distinct commercial antibodies) yields 158,000– 316,000 antibodies estimated to fail independent testing. Using sample counts from publications that used discontinued antibodies without presenting validation data—a restricted subset where the antibody is no longer commercially available—we estimate approximately 4 to 7 million animal samples and 6 to 11 million human tissue samples consumed globally without context-specific validation. These global figures should be read as lower-bound, order-of-magnitude estimates rather than precise counts, dependent on uncertain assumptions and in particular on the extent to which the failure rate observed in the 614-antibody panel generalises to the wider commercial antibody market.

**Table 1.** Summary of the global extrapolation of biological sample waste, showing each step and its input range.

| Step | Basis | Value (range) |
| --- | --- | --- |
| Observed failure rate | YCharOS panel (97/614) | 15.8% |
| Distinct commercial antibodies | CiteAb, de-duplicated | 1–2 million |
| Antibodies likely to fail | 15.8% of the market | 158,000–316,000 |
| Samples per failing antibody | 244 papers ÷ 97 | ~23 animal; ~35 human tissue |
| Global animal samples | 158,000–316,000 × 23 | 4–7 million |
| Global human-tissue samples | 158,000–316,000 × 35 | 6–11 million |

This estimate is intentionally conservative on multiple fronts. It counts only antibodies that failed every application tested, includes only explicitly reported sample numbers from indexed publications, and excludes all unpublished research. Because our literature search was limited to publications available online from December 2000 to October 2025, sample consumption from earlier publications—potentially spanning decades of use for some of these antibodies—is not captured. Importantly, the estimate is based solely on samples consumed using discontinued antibodies, yet 19 of the 35 antibodies in our analysis remain commercially available despite failing standardised validation testing. The samples consumed using these commercially available antibodies—which include 5,847 animal samples and 987 human tissue samples in papers without validation data alone—are excluded from the global projection, even though these antibodies failed the same independent testing as their discontinued counterparts. Critically, the estimate also assumes that the remaining 517 of 614 antibodies in the YCharOS panel—those that passed at least one standardised test—have been used appropriately and produce reliable data in every experimental context. Given that antibody performance is highly context-dependent, varying with application, species, tissue type, and experimental conditions, this assumption is almost certainly too generous. The true scale of biological sample waste from inadequately validated antibodies is therefore likely to be substantially higher than this estimate.

### What are barriers to antibody validation – qualitative themes

Focus group participants described barriers across three categories aligned with the COM-B behavioural model [19]: capability (awareness and knowledge), opportunity (time and resources), and motivation (incentives and habits) (Figure 4A). Illustrative quotes mapped to each theme and subtheme are provided in S1 Table.

**Figure 4.**
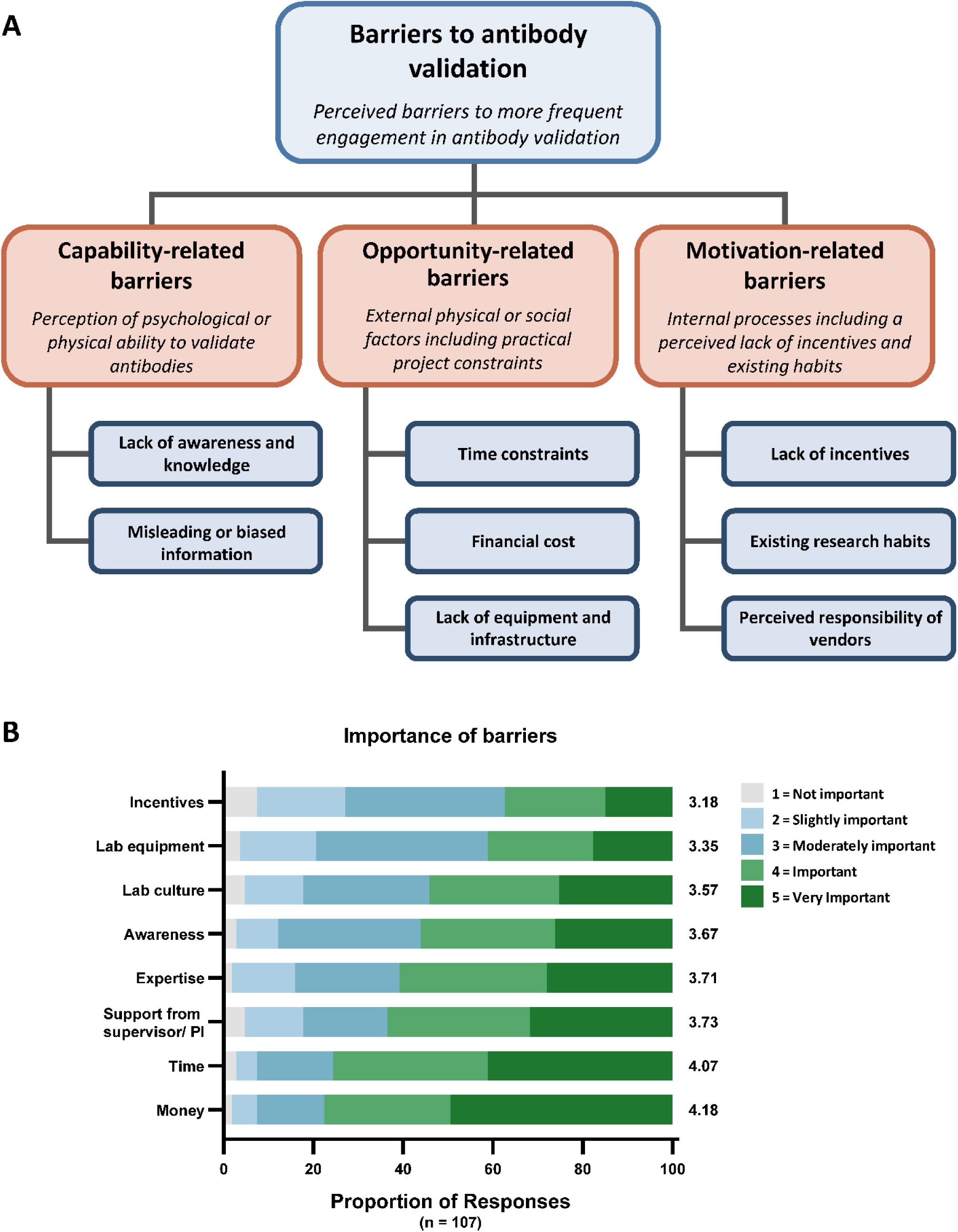
Perceived barriers to antibody validation. (A) Thematic map from focus groups (n=12) categorising barriers using the COM-B model. (B) Survey ratings (n=107) of barrier importance on 5-point Likert scale. Money (77.6%, 83 of 107) and time (75.7%, 81 of 107) were rated as the most important barriers, followed by supervisor support (63.6%, 68 of 107) and expertise (60.7%, 65 of 107), where percentages represent combined ratings of 4 (Important) and 5 (Very Important). Mean Likert ratings are presented to the right of the bars. The data underlying this figure can be found in https://doi.org/10.5281/zenodo.18682587.

Nearly all participants described lack of awareness and knowledge as fundamental barriers, with some having never encountered antibody validation prior to the study. Participants also identified misleading information as a barrier, particularly publication bias favouring positive findings (S1 Table). Time and financial constraints were the most frequently cited opportunity barriers, with one participant characterising antibody validation as an “expensive and time-consuming hobby” (Participant 6). Participants expressed concern that research funders do not adequately account for validation costs (S1 Table).

Some participants questioned whether validation was necessary for widely cited antibodies, while others explained that success with non-validated antibodies provided little incentive to change practice: “You just want nice results. You wanna get a paper? You wanna be successful. So if it’s working, then just go ahead with it” (Participant 12). Participants described research habits that discouraged rigorous validation, such as ignoring unexpected Western blot bands, and some expressed that vendors should bear primary responsibility for validation (S1 Table).

### What are barriers to antibody validation – quantitative survey responses

Survey respondents rated opportunity-related factors of money (77.6%) and time (75.7%) as the most important barriers, followed by supervisor/PI support (63.6%), expertise (60.7%), awareness (56.1%), and lab culture (54.2%). Lack of lab equipment (41.1%) and lack of incentives (37.4%) received lower ratings (Figure 4B).

### What are potential solutions to improve antibody validation – qualitative themes

Focus group participants proposed solutions aligned with behaviour change intervention functions incorporated in the Behaviour Change Wheel [19]: enablement, education, restriction, and incentivisation (Figure 5A). Illustrative quotes mapped to each theme and subtheme are provided in S1 Table.

**Figure 5.**
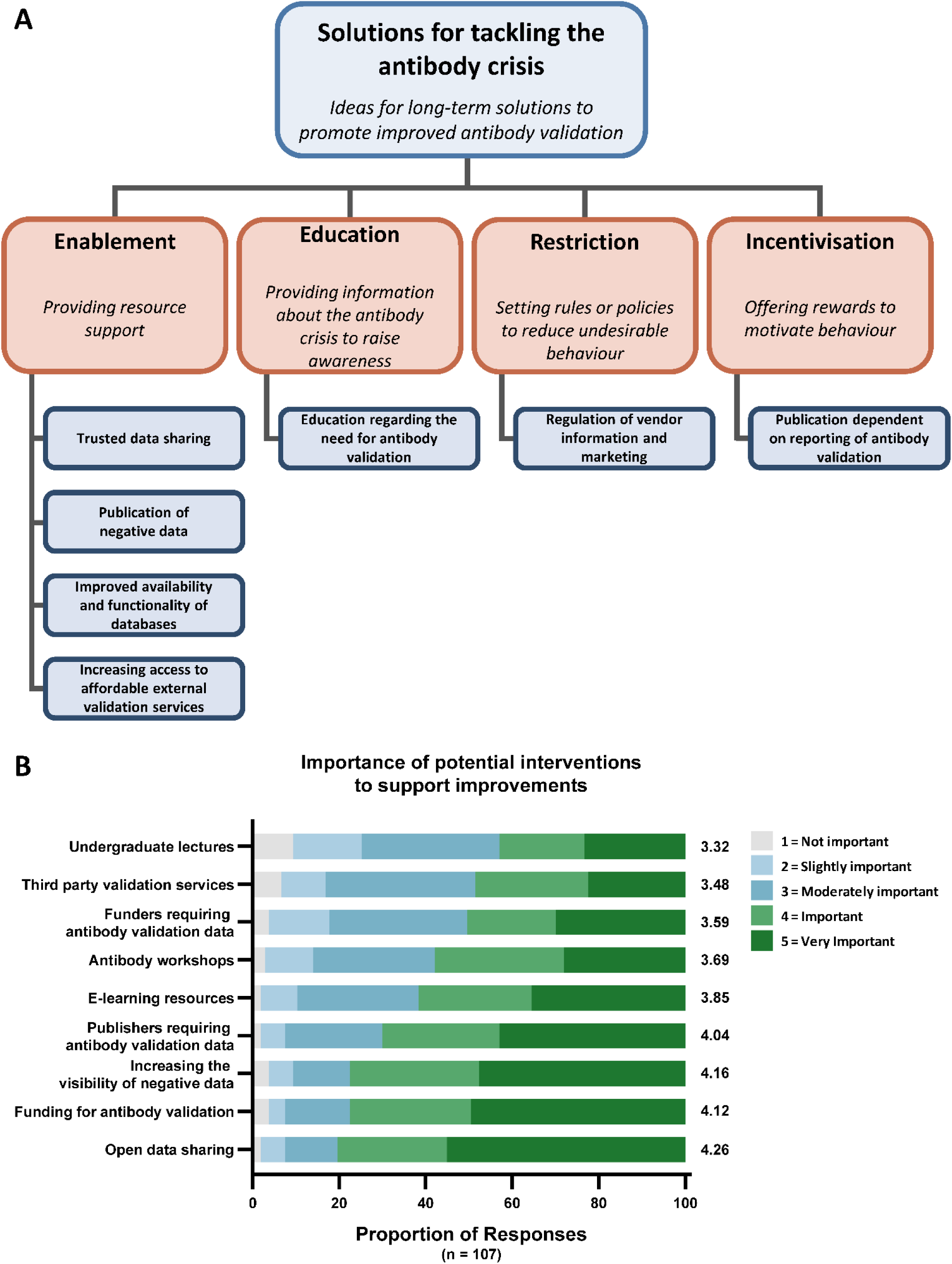
Researcher-proposed solutions to improve antibody validation. (A) Thematic map from focus groups (n=12) organising solutions using Behaviour Change Wheel intervention functions. (B) Survey ratings (n=107) of intervention importance on 5-point Likert scale. Open data sharing received highest support (80.4%, 86 of 107), followed by funding for validation (77.6%, 83 of 107), increasing the visibility of negative data (77.6%, 83 of 107), and publisher requirements (70.1%, 75 of 107), where percentages represent combined ratings of 4 (Important) and 5 (Very Important). Mean Likert ratings are presented to the right of the bars. The data underlying this figure can be found in https://doi.org/10.5281/zenodo.18682587. The most frequently discussed solution was trusted data sharing via neutral websites, particularly including negative results. Participants drew comparisons to consumer review platforms, envisioning a “research hub” where researchers could share antibody outcomes using verified institutional credentials (S1 Table). Participants also suggested better access to external validation services.

Participants emphasised the need for education across all career stages, from undergraduate curricula through to senior researchers, noting that targeting principal investigators could have a multiplier effect on trainees (S1 Table).

In terms of restrictive measures, participants criticised the lack of regulatory oversight for antibody quality and suggested stricter industry standards. Finally, regarding incentivisation, participants proposed that publication requirements for validation evidence could provide clear motivation: “If you’re releasing a paper, there should be a little section proving like how you validated the antibody” (Participant 12; S1 Table).

### What are potential solutions to improve antibody validation – quantitative survey responses

Open data sharing received the highest support (80.4% rating as important or very important), followed by funding for antibody validation (77.6%), increasing visibility of negative data (77.6%), publisher requirements for validation data (70.1%), e-learning resources (61.7%), antibody workshops (57.9%), funders requiring validation data (50.5%), third-party validation services (48.6%), and undergraduate lectures (43.0%) (Figure 5B).

## DISCUSSION

This study set out to understand how researchers select and validate antibodies, quantify the ethical costs of inadequate validation in terms of biological sample waste, and identify barriers and solutions through a behaviourally informed framework. The actionable, stakeholder-endorsed solutions that follow from these findings are set out in our companion Delphi consensus study [20], which we draw on throughout this discussion. This study reveals a substantial gap in context-specific validation data in publications using antibodies with demonstrated performance issues, and provides a lower-bound estimate of the animal lives and human donated biological samples consumed in such research. Drawing on data from antibody end users, it also identifies the behavioural drivers of insufficient validation and the intervention strategies researchers themselves support.

### Drivers of antibody selection and validation assessment

Both focus groups and surveys revealed that antibody purchasing decisions are driven substantially by social factors—previous laboratory use and published data—rather than careful evaluation of performance characterisation data. This reliance on social proof is consistent with broader findings in research decision-making, where heuristics and peer behaviour often substitute for independent evaluation. The dominant form of data evaluated by researchers, Western blot band clarity, while intuitively appealing, is notable given that a single band at the expected molecular weight does not constitute robust evidence of specificity without additional controls [1, 18].

When researchers do evaluate validation data, awareness of formal validation frameworks remains limited. Despite publication in Nature Methods in 2016 and widespread promotion in the scientific literature, only around one-third of surveyed researchers were aware of the IWGAV five-pillar framework [18]. This mirrors the findings of Freedman and colleagues [15], who similarly found limited awareness of validation best practices across a larger sample of 504 researchers in the United States. Whereas that and related work framed the cost of poor antibody performance largely in financial terms, our analysis adds an ethical, sample-based measure: the avoidable use of animal and human tissue. The agreement between two very different samples (senior, United States, 2015; junior, United Kingdom, 2023) on the same barriers and the same awareness gaps suggests these are general features of the field rather than local ones. The persistence of this knowledge gap nearly a decade after the IWGAV recommendations were published underscores the challenge of translating consensus guidelines into laboratory practice [18].

### The validation practice gap

Self-reported validation practices appeared relatively robust, with between 58.9% and 72.0% of survey respondents (depending on whether orthogonal validation is defined strictly or loosely) indicating use of at least one IWGAV-aligned validation method. However, systematic analysis of publications using antibodies with objectively poor performance revealed a stark contrast: only 15.8% of 760 papers presented any validation data. Among the minority presenting validation, genetic strategies dominated, with almost no use of independent antibodies or mass spectrometry approaches [18].

Several mechanisms may explain this substantial gap between self-reported and observed validation. First, the survey asked respondents to indicate which validation methods they had used in their research when using a new antibody for the first time; this may capture lifetime experience rather than routine practice for every antibody. Second, the phrase “positive control cell line” and “negative control cell line” within the survey can be difficult to interpret, and may not always align with the orthogonal pillar within the IWGAV framework as described [18]. Third, social desirability bias may influence survey responses, with researchers overstating adherence to best practices. Fourth, researchers who conduct validation may fail to report it within the paper or supplemental materials. Fifth, the sample of researchers that responded to the survey may not be representative.

A related question is how an antibody can pass a knockout control in one study yet fail knockout-controlled characterisation in another. Examination of the 20 papers reporting a knockout control indicates that the two rarely test the same thing. In 15 of the 20 the knockout material was mouse tissue, whereas the standardised characterisation tested the antibody against the human antigen; and although 14 of the 20 used the antibody for immunohistochemistry or immunocytochemistry, only 4 presented a knockout control in those applications. We also did not evaluate the evidence that antibody signal was abolished in the knockout control in the papers that reported one. Additionally, polyclonal antibodies in particular display lot-to-lot variation (16 of the 20 papers used a polyclonal antibody). It may be that these differences explain the different performances observed.

Regardless of the underlying mechanism, current research cultures and incentive structures inadequately support rigorous antibody validation, with substantial biological material waste, economic costs, and opportunity costs as a consequence.

### Ethical costs: biological sample waste

This study provides, to our knowledge, the first systematic quantification of biological sample waste attributable to the use of poorly performing antibodies without context-specific validation. The central finding is that validation is rare regardless of antibody commercial status (many antibodies have since been discontinued from the market): 84.2% of publications presented no context-specific validation data, and the majority of biological sample consumption occurred in studies not presenting validation data. While Freedman and colleagues estimated approximately $28 billion annually spent on irreproducible preclinical research in the United States, with biological reagents contributing 36.1% [11], the ethical dimension— measured in animal lives and human tissue donations—had remained unquantified. Unlike such financial estimates, which require assumptions about research budget fractions, our biological sample counts provide directly auditable estimates grounded in explicitly reported data — a specificity that strengthens the evidence base for policy interventions.

The strength of these estimates varies by sample type. The human tissue waste estimates rest on the most direct evidence, since the YCharOS pipeline tested antibodies against human antigens [1]—the same species as the tissue samples consumed in the experiments analysed. The inference to animal sample waste involves an additional step, as antibody performance against human antigens does not automatically predict performance against orthologous proteins in other species. However, several lines of convergent evidence support the broader concern. Of the 35 antibodies analysed, 17 were not recommended by their own vendors for use in mouse applications and 16 had been discontinued; collectively, 27 (77.1%) carried at least one of these vendor-level flags, which are independent of YCharOS testing [1]. Combined with the near-total absence of context-specific validation, these data indicate that animal samples were overwhelmingly consumed without any evidence that the antibodies performed adequately in the experimental system used. Antibody performance is highly context-dependent, varying between applications, species, tissue types, and experimental conditions, and arguably the onus is on researchers to demonstrate suitability for their specific use case.

Within this broader pattern, discontinued antibodies represent a particularly acute subset. Research using these antibodies is not only potentially unreliable but also irreproducible, as the reagents can no longer be sourced. Because withdrawal from the market renders these experiments irreproducible irrespective of whether validation was performed but not reported, the waste estimate is insensitive to under-reporting of validation. It is conceivable that an antibody failing standardised testing could perform adequately in a specific experimental context — this is precisely why we examined publications for context-specific validation evidence rather than assuming all use constitutes waste. For the global extrapolation, we centred on antibodies that had also been withdrawn from the market, rendering research using them conclusively irreproducible. Throughout, we reserve the term waste for this most conservative category: antibodies that failed every tested application, were used without reported context-specific validation, and have since been withdrawn from the market. Samples meeting the first two conditions but used with reagents still on sale are described as being at risk of waste rather than demonstrably wasted. Notably, context-specific validation is in many cases inseparable from routine experimental controls — demonstrating that a signal is specific to the intended target is a component of robust experimental design, not a separate exercise undertaken for the antibody’s sake.

Beyond the quantifiable waste lies substantial opportunity cost. These samples, used with appropriately validated reagents, could have contributed to discoveries benefiting patients with severe neurological diseases—the therapeutic area from which most tested antibodies were drawn.

### Barriers and supported interventions

Our findings regarding barriers to validation closely align with those of Freedman and colleagues [15], who surveyed 504 researchers and found that 71% cited time constraints, 49% cost, and 51% indicated that validation delays research. The consistency between studies, despite different populations and methodological approaches, suggests these barriers are robust and cross-cultural.

Mapping barriers onto the COM-B framework [19] reveals failures across all three behavioural domains. Capability barriers include lack of awareness and misleading information from publication bias favouring positive findings. Opportunity barriers—time, cost, and lack of supervisor support—were rated most highly by survey respondents. Motivation barriers include lack of perceived personal benefit from validation when non-validated antibodies produce publishable results, and the belief that widely cited antibodies are already sufficiently validated by the research community.

Despite these barriers, researchers expressed strong support for structural interventions. Open data sharing was rated most important, followed by dedicated funding for validation, increased visibility of negative data, and publisher requirements for validation data. These findings indicate that researchers are receptive to interventions that enable rather than merely restrict behaviour. Future work should engage a broader range of stakeholders—including publishers, funders, manufacturers, and institutional leaders—to identify the most promising and feasible interventions across the research ecosystem. The results of a Delphi consensus study of expert stakeholders to determine evidence-based recommendations for improving antibody validation practice have been reported in our companion paper [20], which sets out a stakeholder-endorsed roadmap of actionable solutions. We emphasise that this is a well-characterised and addressable problem, and that researchers themselves strongly support the relevant interventions.

### Limitations

Several important limitations must be considered. The focus groups included primarily junior researchers with a small sample size (n=12), limiting generalisability to senior investigators. Both focus groups and surveys are subject to selection bias, social desirability bias, and recall bias. The literature analysis can only assess what is reported in publications; validation data that exists, but was not shared or cited, does not necessarily mean validation was not performed.

In interpreting the qualitative and survey data, the relevant standard is information power rather than large-sample representativeness: a narrow question, an established theoretical framework, and well-facilitated discussion reduce the number of participants needed [21]. Neither sample was strategically constructed to mirror the antibody-using community, and both were recruited by opportunity sampling. This biases participation towards researchers already engaged with reproducibility, so our findings are better read as a best case; if these participants show such gaps, the wider community is unlikely to do better. The survey and focus groups also over-represent junior researchers, though these are the people who carry out most experiments and make many day-to-day antibody decisions, so their practice is directly relevant. The perspectives of senior investigators, who may weigh time and cost differently, are under-represented and warrant further study. That said, the larger and more senior, US-based sample of Freedman et al. [15] reported the same leading barriers and the same limited awareness of validation frameworks, suggesting these patterns are not an artefact of our cohort’s junior, UK-weighted composition.

The strongest evidence in our ethical-cost analysis concerns human tissue, which shares the species of the YCharOS antigens; the animal-sample estimates require an additional inferential step and are correspondingly more uncertain. The YCharOS data also derive from cancer cell lines rather than the normal tissues used in much of the surveyed literature. One recent knockout-controlled characterisation speaks to both concerns, although it is limited in scope. In a study of five antibodies against a single target, PPP2R5D [22], the antibodies that passed western blot, immunoprecipitation and immunofluorescence were also the only ones to pass immunohistochemistry, while those that failed these applications were also negative by immunohistochemistry; the passing antibodies further gave the expected knockout-controlled staining in normal mouse brain, lung and muscle. This is consistent with concordance both across applications and between cancer-cell-line and normal-tissue performance, but it is a single target and a small panel and cannot establish that such concordance holds generally. Standardised, knockout-controlled characterisation has to date been established for western blot, immunoprecipitation and immunofluorescence [17], with immunohistochemistry only now being incorporated into such platforms. Systematic cross-application, cross-species and cross-tissue concordance data are correspondingly sparse. We cannot exclude that an individual antibody failing in a cancer line might perform in a particular tissue, or the reverse, and wider cell-line-versus-tissue concordance testing across more targets and tissues would help settle this. These uncertainties do not affect the most conservative estimates, which are restricted to antibodies that failed every application tested, were used without any reported context-specific validation, and have since been withdrawn from the market. Research using such reagents is irreproducible regardless of application or sample type, so the waste calculation stands independently of how far performance generalises across applications.

However, we concentrated our analysis on antibodies with objective performance data demonstrating poor results when tested using standardised protocols—many subsequently removed from the market. For such antibodies, it is reasonable to expect researchers to explicitly demonstrate suitability for their specific use case, as antibody validation is highly context-dependent. Performance can vary between applications, species, tissue types, and experimental conditions. A comparison with publications using antibodies that passed standardised testing could yield different validation rates.

Our biological sample waste estimates represent highly conservative minimum bounds. We included only explicitly reported sample numbers, excluded unpublished research and older research (before December 2000), counted only antibodies failing all tested applications, and relied on conservative market size estimates. Missing data around reagent use means actual waste is likely substantially higher.

The reclassification of antibody commercial status during this study highlights the dynamic nature of the antibody market. Antibodies may be discontinued, reintroduced, or have their recommended applications revised over time, which introduces uncertainty into retrospective analyses. We have reported the status at the time of our analysis.

Finally, our assessment of antibody performance rests on a single standardised dataset. The YCharOS panel provides uniform, knockout-controlled data generated to a published consensus protocol [17], which is what makes comparison across antibodies possible, but it is one source covering 614 antibodies concentrated on neuroscience-relevant targets, and we did not attempt to integrate the substantial body of antibody characterisation published over the preceding four decades. That literature is heterogeneous in method, threshold and reporting, and combining it into a single comparable measure of performance was beyond the scope of this study. Systematic analysis of that wider literature, which machine-assisted methods now make feasible, is an important subject for future enquiry.

## MATERIALS AND METHODS

### Study design

We employed a sequential mixed-methods design to examine how biomedical researchers choose and validate antibodies, and to explore perceived barriers to best practice and potential solutions. The study comprised three components: (1) qualitative focus groups to elicit behavioural themes, (2) a quantitative survey to assess the prevalence of identified behaviours and perceptions across a broader population, and (3) a literature-based analysis estimating frequency of best practice in published manuscripts, and the downstream animal and human sample volume potentially wasted due to poorly characterised antibodies. The focus groups and survey were also used to explore the underlying reasons for lack of validation observed in the literature, and to explore ideas for supporting the uptake of best practices in antibody validation.

The focus group component was designed to explore decision-making processes, perceived capability and infrastructure barriers, and ideas for improving practices. These themes were mapped against theoretical constructs from behaviour change theory, notably the COM-B model and Behaviour Change Wheel [19], to characterise behavioural drivers and intervention opportunities. The survey was then developed to test these themes quantitatively and assess generalisability. Finally, a structured literature analysis sought to evaluate the ethical cost of under-characterised antibodies by estimating animal and human sample use associated with their deployment in published research. The literature analysis was also used to contextualise self-reported behaviours against observed practice.

The study was approved by the University of Leicester Research Ethics Committee (focus group approval reference: 39806-emk12-ls:psychology&visionsciences,schof and survey approval reference: 42401-hsv6-ls:respiratorysciences), and all participants gave written informed consent under GDPR-compliant protocols (focus-group participants via a signed consent form, and survey participants via a mandatory electronic consent item completed before participation). Focus group participants received a £25 Amazon voucher for their time, and survey participants received a £5 Amazon voucher. All participants were offered the option to receive a summary of the study findings.

### Focus groups

We conducted three virtual focus groups between June and July 2023, each comprising four participants (n = 12 total). Participants were UK-based researchers or research students with experience using antibodies in biomedical research. Recruitment used a snowball sampling approach via social media and professional networks. Interested participants were provided with a participant information sheet, gave informed consent in accordance with GDPR, and received a £25 Amazon voucher in appreciation of their time.

Focus groups were conducted via Microsoft Teams and lasted approximately 60–90 minutes. Each session was facilitated by three members of the research team: one led the discussion using a semi-structured topic guide (S1 File), a second displayed relevant visual materials intended to prompt discussion (including Western blot images from commercial vendor sites and a diagram of the IWGAV validation pillars [18]), and a third monitored transcription and group dynamics. Participants were asked about their experiences selecting and using antibodies, the challenges they encountered with validation, and their ideas for improving current practice.

Focus group discussions explored factors influencing antibody selection and validation practices. Participants were asked to reflect on decision-making processes, resource constraints, and common practices they had observed or experienced in their research environments. The study aimed to understand broader psychosocial and systemic barriers rather than audit individual compliance with validation guidelines. Participants were encouraged to speak freely without judgement, but had the right to skip questions and remain silent if preferred. All participants were aware of their right to withdraw from the session at any time.

All sessions were recorded, transcribed verbatim, and de-identified prior to analysis. Transcripts were reviewed and edited by the research team to correct automated transcription errors. A reflexive thematic analysis was conducted following the steps outlined by Braun and Clarke [23], including familiarisation with transcripts, initial coding, generation and refinement of candidate themes, and final thematic mapping. The COM-B model (Capability, Opportunity, Motivation – Behaviour) and Behaviour Change Wheel [19] were used as interpretive frameworks to structure emerging findings. Multiple researchers evaluated the focus group transcripts in full to develop a thematic understanding that was formalised by EK.

### Survey

Following the focus groups, we designed a cross-sectional online survey to assess the generalisability of the themes that emerged. The survey included structured questions on how researchers choose antibodies, their awareness and application of validation strategies, perceived barriers to best practice, and attitudes toward potential solutions. Items were designed to align with behavioural constructs from the COM-B model and the Behaviour Change Wheel [19].

Survey questions were hosted using Microsoft Forms and included multiple choice and 5-point Likert scale formats. Participants were also asked about their experience selecting antibodies, the antibody-based techniques they used, and their awareness or use of key antibody-related resources. The full survey instrument is available in S2 File.

The survey was distributed online in October 2023 via professional networks, institutional mailing lists, and social media using a snowball sampling approach. Eligibility was open to researchers and research students internationally, provided they had experience using antibodies and could confirm affiliation with a recognised academic or industrial research institution. Participants provided informed consent and received a £5 Amazon voucher as a token of appreciation upon completion. Only participants with a confirmed institutional email address (academic or industrial) were included. A total of 107 valid responses were collected.

### Classification of validation approaches from survey responses

To assess alignment between current practice and recommended validation standards, we categorised participants’ self-reported validation methods according to the IWGAV pillar framework [18]. Survey responses for specific validation methods were mapped to IWGAV pillars as follows:

#### Genetic strategies pillar

Participants reporting use of clustered regularly interspaced short palindromic repeats (CRISPR) knockout OR small interfering RNA (siRNA)/short hairpin RNA (shRNA) knockdown were classified as using genetic validation approaches.

#### Tagged expression pillar

Participants reporting comparison of antibody staining between the antibody in question and a protein tag (e.g., his-tag, Green Fluorescent Protein (GFP)) were classified as using tagged expression validation.

#### Independent antibodies pillar

Participants reporting comparison of binding to an independent antibody targeting the same protein were classified as using independent antibody validation.

#### Immunocapture mass spectrometry (MS) pillar

Participants reporting use of MS on samples bound to the antibody were classified as using immunocapture MS validation.

#### Orthogonal validation pillar

This category encompassed two distinct approaches. First, participants reporting use of antibody-independent methods (e.g., transcriptomics, proteomics) for triangulation were classified as using orthogonal validation. Second, we classified participants who reported using BOTH positive control cell lines (known to express the target) AND negative control cell lines (known not to express the target) as employing a form of sample-based orthogonal validation, as this approach requires independent knowledge of target expression in the cell lines used and enables comparison of antibody signal against known expression status. Participants reporting use of only positive OR only negative control cell lines were not classified under this category, as comparison between both is necessary for this validation strategy.

Participants were classified as IWGAV-aligned if they reported using at least one method corresponding to any of the five categories above. Participants reporting no IWGAV-aligned methods but using at least one technical control (fluorescence minus one, isotype control, secondary-only control, blocking peptide, or single positive/negative control lines) were classified as technical controls only. The remaining participants were classified as reporting no validation.

Descriptive statistics were used to summarise the data, and responses were visualised using stacked bar charts. Survey findings were used to corroborate or contrast the qualitative themes identified in the focus groups.

### Literature analysis to assess frequency of observed antibody validation and ethical costs of insufficient validation

We focused on 35 commercial antibodies that were systematically tested by the Antibody Characterisation by Open Science (YCharOS) consortium using standardised, industry-endorsed protocols [17,24] and were found to perform poorly across multiple applications. We identified publications citing these antibodies and assessed whether validation data were presented.

We identified a subset of commercial antibodies evaluated in [1]. To focus the literature search on antibodies that critically require context-specific validation, we selected antibodies found to perform poorly in multiple applications when assessed according to industry-endorsed, standardised protocols [17]. From 97 commercial antibodies that failed YCharOS validation testing in all applications (out of 614), we excluded three antibodies primarily recommended for neutralising assays, as these do not lend themselves to validation assessment using the IWGAV five-pillar framework. Literature searches using CiteAb and Google Scholar identified publications for 35 of the remaining 94 antibodies by matching vendor catalogue numbers; 59 antibodies had no associated publications.

Each identified publication was reviewed to determine: (1) whether any antibody validation data were presented according to any of the IWGAV [18] five pillars; (2) which experimental methods used the antibody in question; and (3) the number of human or animal biological samples involved in those experiments.

Only sample numbers explicitly reported in figure legends or methods sections were included. No assumptions or extrapolations were made based on standard practice or technique type. If sample counts could not be determined from the publication, no estimate was recorded for that study. The analysis excluded unpublished data and publications not indexed or matched via CiteAb.

The resulting values were aggregated to produce a minimum-bound estimate of the human and animal sample burden per antibody and across the reviewed set.

### Estimation of global biological sample waste from insufficient antibody validation

To provide an order-of-magnitude estimate of the ethical costs of insufficient antibody validation, we extrapolated our findings to the broader commercial antibody market.

#### Step 1: Per-antibody sample consumption rates

We calculated per-failing-antibody consumption rates using samples from publications employing discontinued antibodies that presented no validation data (n=244 papers: 2,217 animal samples, 3,437 human tissue samples, 599 human primary cell samples, 153 cell lines), divided by 97 (total antibodies failing all tests), yielding approximately 23 animals, 35 human tissue samples, 6 human primary cell samples, and 2 cell lines per failing antibody. This approach is deliberately conservative: it restricts the numerator to samples from discontinued antibodies without validation—a subset where the manufacturer has removed the product from the market—but divides by all 97 failing antibodies, deflating the per-antibody rate.

#### Step 2: Market size estimation

The CiteAb antibody database indexes approximately 8 million antibody products from over 300 suppliers [25], though this includes substantial duplication as the same clone is often sold by multiple distributors. Individual large vendors report catalogues of >120,000 products (e.g., Abcam [26]), and industry analyses describe “several million distinct products” globally [27]. Accounting for cross-vendor duplication, we adopted a conservative estimate of 1–2 million distinct commercial research antibodies.

#### Step 3: Extrapolation calculation

We applied the failure rate from our 614-antibody panel (97/614 = 15.8%) to the market size estimate, yielding 158,000–316,000 potentially failing antibodies. Multiplying by per-antibody consumption rates from Step 1 gives an estimated 4–7 million animal samples and 6–11 million human tissue samples consumed globally without context-specific validation.

This extrapolation is designed as a lower bound. It excludes samples consumed using the 19 antibodies that remain commercially available despite failing standardised testing, publications reporting any validation data (even if incomplete or conducted in a different context), unpublished research, and studies where sample numbers were not explicitly reported. Our literature search was also limited to publications available online from December 2000 to October 2025, meaning earlier publications are not captured. Additional assumptions include: (1) only antibodies failing all tested applications are counted; (2) only publications indexed by CiteAb and Google Scholar with matching catalogue numbers are included; (3) our 614-antibody panel is assumed to exhibit similar failure rates to the broader market; and (4) the true number of distinct commercial antibodies is unknown. It should also be noted that antibodies may be discontinued for a range of commercial or technical reasons, not exclusively due to quality concerns.

#### Species considerations

The YCharOS standardised testing was conducted against human antigens. Our human tissue waste estimates therefore represent the most directly supported inference. The inference to animal waste is indirect but supported by convergent evidence: 17 of 35 antibodies (48.6%) were not recommended for mouse by their vendors and 16 (45.7%) were discontinued, with 27 (77.1%) carrying at least one of these flags—determinations independent of YCharOS testing. Combined with the finding that 84.2% of papers presented no context-specific validation, the animal waste estimates reflect a well-founded concern, though they carry additional uncertainty relative to the human tissue estimates.

## DATA AND METHODS AVAILABILITY

The focus group guide, survey instrument, survey responses (n=107), complete literature analysis data (97 antibodies, 785 publications, sample counts, validation assessments, and publication metadata) are openly available in Zenodo at DOI: https://doi.org/10.5281/zenodo.18682587. This deposition contains the individual observations underlying every figure panel. Figures were plotted in GraphPad Prism directly from these data; no custom code was generated during this study. Focus group transcripts contain potentially identifying information and cannot be made publicly available in accordance with participant consent and ethics approval (University of Leicester Research Ethics Committee reference: 39806-emk12-ls:psychology&visionsciences,schof). De-identified representative quotes are included in S1 Table. S2 and S3 Tables provide summary statistics from the literature analysis.

## SUPPORTING INFORMATION

**S1 File.** Focus group materials. Complete focus group protocol including participant information sheet, consent form, pre-focus group questionnaire, topic guide, and example antibody data sheets used for discussion prompts.

**S2 File.** Survey instrument. Complete survey questions administered via Microsoft Forms, including Likert-scale items on antibody selection factors, validation practices, barriers, and proposed solutions.

**S1 Table.** Representative quotes from focus group discussions (n=12 participants across 3 sessions) organised by results section and theme. Quotes are mapped to the themes and subthemes identified through reflexive thematic analysis, providing the evidential basis for the qualitative findings reported in the main text.

**S2 Table.** Biological sample counts by antibody and validation status. Complete breakdown of animal samples, human tissue samples, human primary cell samples, and cell lines identified across 785 publications, stratified by antibody commercial status (discontinued/continued) and validation evidence (present/absent).

**S3 Table.** Validation methods observed in publications. Frequency of IWGAV-aligned validation approaches (genetic strategies, tagged expression, independent antibodies, immunocapture MS, orthogonal validation) across 120 publications presenting validation data.

## ACKNOWLEDGEMENTS

The YCharOS consortium manufacturers and the rest of the YCharOS consortium for providing materials for testing and sharing the open data.

## ABBREVIATIONS

COM-B: Capability, Opportunity, Motivation–Behaviour (model)
CRISPR: clustered regularly interspaced short palindromic repeats
GFP: Green Fluorescent Protein
ICC/IF: immunocytochemistry/immunofluorescence
IWGAV: International Working Group for Antibody Validation
MS: mass spectrometry
shRNA: short hairpin RNA
siRNA: small interfering RNA
YCharOS: Antibody Characterisation by Open Science (antibody validation consortium)

