## Supplementary material for "Inadequate antibody validation places substantial numbers of animal and human tissue samples at risk of waste": S1 Table

**S1 Table. Illustrative focus group quotes supporting qualitative themes.**

Representative quotes from focus group discussions (n=12 participants across 3 sessions) organised by results section and theme. Quotes are presented verbatim with minor edits for readability indicated by square brackets. Participant identifiers (P1–P12) are consistent across sections. Quotes are mapped to the themes and subthemes identified through reflexive thematic analysis, providing the evidential basis for the qualitative findings reported in the main text. Full transcripts are not publicly available in accordance with participant consent and ethics approval.

| Results Section | Theme / Subtheme | Illustrative Quote |
|---|---|---|
| Antibody selection | Social and reputational factors | *"[...] similar things that already been mentioned about the reputability of the journal and the authors... might see if it's been cited by other people"* (P1) |
| | | *"if there's more references on one than the other, that tends to influence my choice quite a lot"* (P4) |
| | | *"The first thing I do actually is probably ask around in the lab if anyone else's sort of tried to do what I'm about to do, they've used an antibody [...] And then I'd probably just go with what they've gone with"* (P7) |
| | | *"I've just been told what to pick. [...] I've never even heard of the word validating antibodies until [...] this meeting"* (P12) |
| | Convenience and pragmatic concerns | *"once you've got one that works, you stick with it"* (P3) |
| | | *"They tend [to be] actually more expensive, but at least I have a degree of confidence that they will be staining what I need them to stain"* (P6) |
| | Engagement with validation data | *"[...] maybe see what data the supplier has. So, if they've got some example Western blots... Looking at some of the quality of the bands that have produced, is it nice and clean or is it more of a broad band"* (P7) |
| Barriers to validation | Capability: Awareness | *"Nope, I've got no clue how they chose it, and I'm guessing for my PhD, I know that the answer has been shown to work..."* (P12) |
| | | *"I didn't know that was such a problem"* (P2) |
| | Capability: Misleading information | *"studies that make it to the literature only gonna be presenting positive findings were the antibodies worked really nicely. Whereas there may be [...] thousands of [...] other people who've tried to use the antibody and it hasn't worked"* (P7) |
| | Opportunity: Time and cost | *"For you to do a proper validation of an antibody that can take months or years, especially if there's a lot of antibodies for [a] specific protein in your [...] field of work"* (P6) |
| | | *"Sponsors or grants are not gonna want you to spend your... that kind of money on a wild goose chase, to be fair"* (P6) |
| Barriers to validation | Motivation: Lack of incentive | *"a lot of the antibodies that I use are... extremely well used and validated and used by thousands of immunologists. And they are kind of just the standard that we use"* (P5) |
| | | *"You just want nice results. You wanna get a paper? You wanna be successful. So if it's working, then just go ahead with it and that we don't need to validate it"* (P12) |
| | Motivation: | *"Somewhere else on the blot you see something else and you're a bit like...* |

| Results Section | Theme / Subtheme | Illustrative Quote |
|---|---|---|
| | Research habits | *Absolutely no idea what that is. It's not supposed to be there [...] no one will ask a question"* (P4) |
| | Motivation: Vendor responsibility | *"maybe the company's not taking full responsibility of actually checking the antibodies before pushing out to market, maybe cutting corners [...] just so they can get more of a profit"* (P5) |
| Proposed solutions | Enablement: Data sharing | *"I would really like to see that someone has tried this antibody and it hasn't worked"* (P1) |
| | | *"Just like maybe like [...] any of the food apps [...] the researchers, who had a login information with their [...] university address [...] can log on and then publish their [...] results. So if they have used this, tried this antibody: What was outcome? [...] So, it's basically like a research hub [...] helping each other out"* (P8) |
| | Enablement: External services | *"I think we could reduce this cost of validating antibody by using external provider or using the university facility"* (P9) |
| | Education | *"Teaching it at an undergraduate level would probably be quite helpful, because then you'd get people going into research already knowing that this is quite an important thing that you have to do"* (P10) |
| | | *"Would it make more sense to advertise to those that are higher up? So maybe like PI's, cause they have more influence over more people. So, if you get them on board, then they can influence [...] PhD students"* (P10) |
| | Restriction | *"would want the company that manufactured the antibody to [...] have to do some sort of test and show that data"* (P2) |
| | | *"there's no one who is [...] looking for that for it. There's no authority"* (P8) |
| | Incentivisation | *"If you're releasing a paper, there should be a little section proving like how you validated the antibody before going on to your results. Cause then you [...] prove [...] it's real"* (P12) |
