## Supplementary material for "Inadequate antibody validation places substantial numbers of animal and human tissue samples at risk of waste": S2 Table

### S2 Table. Biological sample usage by antibody commercial status and validation evidence

| Sample Category | Papers (n) | Animals‡ | Human Tissue | Human Primary Cells | Cell Lines |
|---|---|---|---|---|---|
| **ALL PAPERS*** | 766 | 9,443 | 5,015 | 1,215 | 569 |
| **By antibody commercial status:** | | | | | |
| Discontinued (16 antibodies) | 307 | 3,036 | 3,910 | 729 | 236 |
| Continued (19 antibodies) | 459 | 6,407 | 1,105 | 486 | 333 |
| **By validation evidence†:** | | | | | |
| **WITHOUT validation** | 640 | 8,064 | 4,424 | 1,081 | 400 |
| Using discontinued antibodies | 244 | 2,217 | 3,437 | 599 | 153 |
| Using continued antibodies | 396 | 5,847 | 987 | 482 | 247 |
| **WITH validation** | 120 | 1,339 | 513 | 134 | 168 |
| Using discontinued antibodies | 60 | 815 | 421 | 130 | 83 |
| Using continued antibodies | 60 | 524 | 92 | 4 | 85 |

**Notes:**

* Sample numbers were extractable from 766 of 785 publications analysed

† Validation status and sample numbers could be determined for 760 papers

‡ Animal samples include mouse, rat, and other species combined
