## Supplementary material for "Inadequate antibody validation places substantial numbers of animal and human tissue samples at risk of waste": S3 Table

### S3 Table. Detailed biological sample usage by technique, antibody status, and validation evidence

#### ANIMAL SAMPLES

##### Mouse/Rat samples

| Category | Papers (n) | Total | WB | IHC/IHC-IF | ICC | FC |
|---|---|---|---|---|---|---|
| All papers (sample numbers identified) | 766 | 8,533 | 7,145 | 1,828 | 477 | 59 |
| Discontinued antibodies (16) | 307 | 2,856 | 2,391 | 365 | 160 | 0 |
| Continued antibodies (19) | 459 | 5,677 | 4,754 | 1,463 | 317 | 59 |
| **Papers with validation & sample data** | 760 | 8,493 | 7,109 | 1,824 | 477 | 59 |
| **WITHOUT validation** | 640 | 7,253 | 6,021 | 1,580 | 359 | 59 |
| Discontinued antibodies | 244 | 2,057 | 1,735 | 256 | 66 | 0 |
| Continued antibodies | 396 | 5,196 | 4,286 | 1,324 | 293 | 59 |
| **WITH validation** | 120 | 1,240 | 1,088 | 244 | 118 | 0 |
| Discontinued antibodies | 60 | 795 | 656 | 105 | 94 | 0 |
| Continued antibodies | 60 | 445 | 432 | 139 | 24 | 0 |

##### Other species samples

| Category | Papers (n) | Total | WB | IHC | ICC |
|---|---|---|---|---|---|
| All papers (sample numbers identified) | 766 | 910 | 726 | 256 | 132 |
| Discontinued antibodies (16) | 307 | 180 | 176 | 0 | 24 |
| Continued antibodies (19) | 459 | 730 | 550 | 256 | 108 |
| **Papers with validation & sample data** | 760 | 910 | 726 | 256 | 132 |
| **WITHOUT validation** | 640 | 811 | 627 | 256 | 132 |
| Discontinued antibodies | 244 | 160 | 156 | 0 | 24 |
| Continued antibodies | 396 | 651 | 471 | 256 | 108 |
| **WITH validation** | 120 | 99 | 99 | 0 | 0 |
| Discontinued antibodies | 60 | 20 | 20 | 0 | 0 |
| Continued antibodies | 60 | 79 | 79 | 0 | 0 |

### HUMAN SAMPLES

#### Human tissue samples

| Category | Papers (n) | Total | WB | IHC | FC |
|---|---|---|---|---|---|
| All papers (sample numbers identified) | 766 | 5,015 | 1,024 | 4,120 | 19 |
| Discontinued antibodies (16) | 307 | 3,910 | 864 | 3,194 | 0 |
| Continued antibodies (19) | 459 | 1,105 | 160 | 926 | 19 |
| **Papers with validation & sample data** | 760 | 4,937 | 1,024 | 4,042 | 19 |
| **WITHOUT validation** | 640 | 4,424 | 859 | 3,654 | 19 |
| Discontinued antibodies | 244 | 3,437 | 724 | 2,821 | 0 |
| Continued antibodies | 396 | 987 | 135 | 833 | 19 |
| **WITH validation** | 120 | 513 | 165 | 388 | 0 |
| Discontinued antibodies | 60 | 421 | 140 | 321 | 0 |
| Continued antibodies | 60 | 92 | 25 | 67 | 0 |

#### Human primary cells/serum samples

| Category | Papers (n) | Total | WB | FC | ICC | ELISA |
|---|---|---|---|---|---|---|
| All papers (sample numbers identified) | 766 | 1,215 | 665 | 366 | 76 | 123 |
| Discontinued antibodies (16) | 307 | 729 | 403 | 224 | 57 | 57 |
| Continued antibodies (19) | 459 | 486 | 262 | 142 | 19 | 66 |
| **Papers with validation & sample data** | 760 | 1,215 | 665 | 366 | 76 | 123 |
| **WITHOUT validation** | 640 | 1,081 | 600 | 366 | 52 | 66 |
| Discontinued antibodies | 244 | 599 | 342 | 224 | 33 | 0 |
| Continued antibodies | 396 | 482 | 258 | 142 | 19 | 66 |
| **WITH validation** | 120 | 134 | 65 | 0 | 24 | 57 |
| Discontinued antibodies | 60 | 130 | 61 | 0 | 24 | 57 |
| Continued antibodies | 60 | 4 | 4 | 0 | 0 | 0 |

### CELL LINES

#### Human cell line samples

| Category | Papers (n) | Total | WB | FC | ICC |
|---|---|---|---|---|---|
| All papers (sample numbers identified) | 766 | 569 | 523 | 5 | 120 |
| Discontinued antibodies (16) | 307 | 236 | 216 | 1 | 57 |
| Continued antibodies (19) | 459 | 333 | 307 | 4 | 63 |
| **Papers with validation & sample data** | 760 | 568 | 522 | 5 | 119 |
| **WITHOUT validation** | 640 | 400 | 364 | 3 | 76 |
| Discontinued antibodies | 244 | 153 | 142 | 0 | 29 |
| Continued antibodies | 396 | 247 | 222 | 3 | 47 |
| **WITH validation** | 120 | 168 | 158 | 2 | 43 |
| Discontinued antibodies | 60 | 83 | 74 | 1 | 28 |
| Continued antibodies | 60 | 85 | 84 | 1 | 15 |

### VALIDATION METHODS IN PAPERS WITH VALIDATION DATA

Total papers where validation status could be determined: 760 (of 766 papers with extractable sample numbers)

Papers with validation data: 120 (60 used discontinued antibodies)

Papers without validation data: 640 (244 used discontinued antibodies)

**Types of validation used in 120 papers with validation data:**

| Validation method | Number of papers | Notes |
|---|---|---|
| Genetic validation: knockout | 20 | CRISPR or other knockout control. The definitive genetic control |
| Genetic validation: knockdown | 83 | siRNA/shRNA knockdown without an accompanying knockout control. A further 6 papers reported both knockdown and knockout and are counted in the knockout row above; 89 papers reported knockdown in total |
| Overexpression/tagged expression | 40 | Tagged protein comparison |
| Independent antibodies | 1 | Comparison with independent antibody |
| Mass spectrometry | 0 | Immunocapture MS validation |
| Orthogonal validation | | Unable to confidently identify |

Note: Some papers used more than one validation strategy, so counts do not sum to 120. Knockout and knockdown together account for 103 papers reporting a genetic validation strategy.

Abbreviations: WB = Western blot; IHC = Immunohistochemistry; IHC-IF = Immunohistochemistry-Immunofluorescence; ICC = Immunocytochemistry; FC = Flow cytometry; ELISA = Enzyme-linked immunosorbent assay
