## Supplementary material for "Inadequate antibody validation places substantial numbers of animal and human tissue samples at risk of waste": S1 File

#### Appendix 1: Study Advertisement

### Do you use antibodies in your research?

Tell us about your experience of choosing, buying and using antibodies in our online focus group discussion with other researchers.

#### You will receive £25 for participation!

Any UK-based researcher or research student that uses/has used antibodies is eligible to participate

For more information or to sign up, please contact:

Example antibody image removed (manufacturer-supplied; licence unverified)

Example antibody image removed (manufacturer-supplied; licence unverified)

Example antibody image removed (manufacturer-supplied; licence unverified)

#### Appendix 2: Participant Information Sheet

University of Leicester
logo removed

**INFORMATION LETTER**

**Focus group to explore how researchers/ post-graduate research students choose and use antibodies.**

**Invitation**

Our names are Dr Harvinder Virk and Dr Eva Krockow from the University of Leicester in the UK. We would very much like to invite you to take part in this study. Before you decide, it is important for you to understand why the research is being done and what it will involve. Please take as much time as you need to read the following information carefully. You can discuss it with others if you wish. Please feel free to contact us, if there is anything that is not clear, or if you would need more information on the study before deciding to participate.

**The purpose of this study**

This study is part of a larger project that aims to give researchers new tools to help them choose and use antibodies in their research. The aim of this focus group is to understand how researchers choose antibodies to use in their research, how they use them, and what their research experiences are with antibodies more generally. If you have had any difficulties with choosing or using antibodies, we would like to explore your ideas for how to overcome these.

**Why have I been invited to participate?**

You have been invited to participate because you have identified as someone who carries out research involving the use of antibodies, and we are interested to know your views about choosing and using antibodies in your research.

**Do you have to take part?**

No. It is completely up to you whether to participate in this study or not. If you decide to participate in this research study, please read and then sign the informed consent document and complete the short pre-study questionnaire. You will also be offered the chance to receive a brief report of our findings, so that you can see how your data has been used. As a thank you for your participation and time we would like to offer you a £25 Amazon voucher.

**What you have to do**

You will attend a 60-90-minute focus group meeting online via Microsoft Teams. You will be joined by up to five other research participants and asked questions as a group about how you choose, use and buy antibodies. We will also ask you to explain which antibody you would choose in a hypothetical decision scenario about antibodies. You do not have to answer each question if you prefer not to. There are no right or wrong answers. We are simply interested in the different approaches researchers use, including their ideas about how to improve things for researchers. After participation, you will be rewarded with a £25 Amazon voucher for your time.

**What are the possible disadvantages and risks of taking part?**

There are very few possible disadvantages or risks of taking part. One disadvantage is that you will have to give up approximately 60-90 minutes of your free time.

**What are the possible benefits of taking part?**

You will receive a remuneration for your time in form of a £25 Amazon voucher. Taking part in this research study could also potentially benefit other researchers by providing insights into factors that shape antibody choices. Ultimately, this study aims to identify strategies to improve the process of antibody choice and use for the benefit of researchers and their research.

**What data will you collect about me?**

We will minimise the data collected about you. We use your name and contact information to organise the focus groups and ensure later payment, but will delete these data immediately after data collection is completed. We will collect some demographic data to check for sample diversity. Additionally, we will ask a few questions about your previous research experience with antibodies, because this is likely to affect your attitudes on the topic.

**Will your taking part in this study be kept confidential?**

All of the information collected during this research study will be kept strictly confidential. No names or identifiable information will be collected, and all collected data will be securely stored. However, it will not be possible to keep you anonymous during the focus group discussion, which will be facilitated by two researchers from our team, and involve up to 5 other participants. We would ask those who take part in the discussion to keep any sensitive content confidential, however we do not have any legal contract with participants to enforce this. However, we do not expect the focus group to discuss very sensitive topics and participants will not be asked personal questions. Furthermore, participants are not obliged to answer all questions addressed as part of the focus group.

**How will you look after the data you collect about me?**

We need to ensure that you understand what will happen to data we collect about you as well as your legal rights. This document is accompanied with a separate Privacy Notice providing further details, which will follow on the next page.

At all times this research study will comply with the General Data Protection Regulations (GDPR, 2018) approved by the EU parliament on 14 April 2016 and passing into UK law with effect from 25 May 2018.'

If you wish to withdraw from the study, you may do so at any point up to 2 weeks following participation in the focus group. This will result in the transcript of your data being removed from the study. However, because this is a focus group discussion, it will not be possible to remove the influence your participation will have had on the other participants and their responses.

**What will happen to the results of the study?**

The information collected will be analysed and written up, and the findings may be published in peer reviewed scientific journals or presented at academic conferences. The results will also inform how to produce new tools to better support researchers in antibody selection and use.

**What should I do if I want to take part?**

You will be asked to complete an Informed Consent Form and to opt-in to a variety of research options by ticking the Yes or No box. This will confirm you understand how your data will be processed, protected and reviewed for research purposes.

**Who is organising and funding the research project?**

This study is being conducted by staff from the College of Life Sciences at the University of Leicester. The principal investigators are Dr Harvinder Virk from the Department of Respiratory Sciences and Dr Eva Krockow from the School of Psychology and Vision Sciences. We are funded by the Leicester Institute of Advanced Studies and we have no commercial interests in companies that make or sell antibodies.

**What if something goes wrong?**

In the very unlikely event of you being harmed by taking part in this research project, there are no special compensation arrangements. If you are harmed due to someone's negligence, then you may have grounds for legal action but you may have to pay for it.

**Who has reviewed the research project?**

This project has been approved by the University of Leicester Research Ethics Committee. If you require further information about the study, please email Dr Eva Krockow. If you have any comments about the conduct of this study, please contact the Chair of the Ethics Committee, Dr Ainslea Cross.

Thank you

#### Appendix 3: Privacy Notice

University of Leicester
logo removed

### Privacy Notice for Research Participants

#### Focus group to explore how researchers/ research students choose and use antibodies.

Dr Harvinder Virk and Dr Eva Krockow

This Privacy Notice provides information about how the University of Leicester collects and uses your personal information when you take part in this research projects.
Please also refer to the Participant Information Sheet given to you for further details about the research project, what information will be collected about you, and how it will be used.
The University of Leicester will usually be the *Data Controller* of any data that you supply for this research. This means that we are responsible for looking after your information and using it properly. This means that the University will make the decisions on how your data is used and for what reasons. The exception to this is joint research projects, if this is applicable you will be informed on the Participant Information Sheet as to the other partner institution(s) who will also have responsibilities for looking after your information. You can access more information on this via the University's Information Assurance Services:

Information Assurance Services
University of Leicester
University Road
Leicester
LE1 7RH
T: +44 (0)116 229 7945
E:
W: https://www2.le.ac.uk/offices/ias

**Why do we need your data?**
Your data will improve our understanding of how researchers choose and use antibodies in their research. We wish to understand any difficulties that researchers might currently experience and their ideas for overcoming these.

**University of Leicester's legal basis for collecting this data is:**
Processing is necessary for the performance of a task in the public interest such as research.

If the university asks you for sensitive data such as; your racial or ethnic origin, political opinions, religious or philosophical beliefs, trade-union membership, data concerning health or sexual life, genetic/biometric data or criminal records, the University of Leicester will use these data because processing is necessary for scientific or research in the public interest.

**What type of data will the University of Leicester use?**
The focus group will generate audio and video data through recordings of the focus groups. Once data collection is completed, the data will be transcribed verbatim and any identifiable information (e.g. names or places names) will be removed. Then, the original recordings will be recorded and the transcribed, anonymised data will be stored in MS Word documents.

**Who will the University of Leicester share your data with?**
Only the project leads (Dr Virk and Dr Krockow) and immediate project researchers Dominic Ruddy will have access to the raw identifiable data (i.e. the audio and video recordings) before they are deleted. The anonymised data transcripts may be shared with collaborating researchers from the University of Leicester or other universities

The anonymised, transcribed data may also be shared via the University of Leicester research repository *Figshare*, which is used to make raw research data available to other researchers.

**Will the University of Leicester transfer my data outside of the UK?**
No.

**What rights do I have regarding my data held by the University of Leicester?**
Your normal rights under the Data Protection Act and the General Data Protection Regulation apply. However, we need to manage your records in specific ways for the research project to be reliable. This means that we will not [always] be able to let you see or change the data we hold about you.

You are free to withdraw at any time during the period of the study, and up until two weeks after participation by emailing the researchers. This will result in the transcript of your data being removed from the study. However, because this is a focus group discussion, it will not be possible to remove the influence your participation will have had on the other participants and their responses.

**Where did the University of Leicester source my data from?**
All data are collected directly from research participants.

**Are there any consequences of not providing the requested data?**
There are no consequences of not providing data for this research. It is purely voluntary.

**Will there be any automated decision making using my data?**
There will be no use of automated decision making in scope of UK Data Protection and Privacy legislation.

**How long will the University of Leicester keep my data?**

In line with the law, we will only keep your data for as long as we need to so that we can fulfil our research objectives.

We will keep your personal data, such as your name and email address until we have completed all the actions that require us to hold them, for example sending you a copy of the results of the study if you have requested this, and then the data will be destroyed. This will take no longer than 12 months from when the study ends.

We will keep the original audio and visual materials until we have completed data collection and transcribed the data verbatim. Then, the recordings will be destroyed. This will take no longer than 12 months from when the study ends.

Anonymised, transcribed data will be kept in electronic form indefinitely and may be shared with other researchers upon request or via the University of Leicester research repository *Figshare.*

**Who can I contact if I have concerns?**
In the event of any questions about the research project, please contact the researchers in the first instance.
Dr Harvinder Virk:
Dr Eva Krockow:

If you have any concerns about the way in which the research project has been conducted, please contact the **Chair of the University Research Ethics Committee** at.

The University of Leicester Data Protection Officer is:

Data Protection Officer
University of Leicester,
University Road, Leicester, LE1 7RH
0116 229 7640


For further details about information security, please contact the Information Assurance Services team: https://le.ac.uk/ias.

### Appendix 4: Participant Consent Form

University of Leicester
logo removed

#### CONSENT FORM

**Full title of Project: Focus group to explore how researchers choose and use antibodies.**

**Name, position and contact details of Researcher:**

Dr Harvinder Virk, Clinical Lecturer,

Dr Eva Krockow, Lecturer in Psychology,

Please **initial** box

1. I confirm that I have read and understand the participant information sheet (Version 2.0, 15.03.2023) for the above study and have had the opportunity to ask questions. ☐
2. I understand that my participation is voluntary and that I am free to withdraw at any time during the period of the study, and up until two weeks after participation by emailing the researcher. ☐
3. I understand that at all times this research project will comply with the *General Data Protection Regulations (GDPR, 2018)* approved by the EU parliament on 14 April 2016 and passing into UK law effective from 25 May 2018 and that if I have any concerns how I contact the University of Leicester to raise these. ☐
4. I understand that the focus will be audio-recorded and video-recorded. ☐
5. I agree to take part in the above research project. ☐

Please **initial** box

| | Yes | No |
|---|---|---|
| 6. I agree to the use of anonymised quotes in publications. | ☐ | ☐ |

7. I agree that anonymised information, gathered about me for this research project may be stored in the University of Leicester research repository Figshare, which is used to make raw research data available to other researchers. ☐ ☐

8. I agree that anonymised data collected for this research project may be used in future research. ☐ ☐

9. I wish to receive a copy of the results of this research project, and I agree for my contact details to be retained and used for this purpose. ☐ ☐

| Name of Participant | Date | Signature |
|---|---|---|

Ethics application number: 39806-emk12-ls:psychology&visionsciences,schof

#### Appendix 4: Pre-Focus Group Questionnaire

##### Questionnaire

If you consent to taking part in this study, please complete the following questions and return the questionnaire alongside the consent form. Please note that the data provided here will be anonymised and kept separately from your personal information (name and contact details).

1. Please indicate your age in years: ___________

2. Please indicate your sex:
- Male
- Female
- Other
- Prefer Not to Say

3. Highest (completed) education Level:
- College sixth form graduate or equivalent
- Bachelor's degree or equivalent
- Master's degree or equivalent
- PhD or equivalent

4. What statement best describes your research experience level?
- Undergraduate level
- Masters level
- PhD student
- Post-Doc/ Early career researcher
- Experienced Post-Doc
- Independent investigator
- Other (please specify)_______________________________

5. Please state which research institution(s) you work/ or have worked in in the past 5 years:
_________________________________________________________________________
_________________________________________________________________________

6. Who funds, or has funded, your research work involving antibodies (tick all that apply)

- Government/ Research Council
- Commercial
- Charity
- Other

7. Have you ever accessed or used any of the following antibody related resources?

CiteAb

Antibodypedia

ProteinAtlas

YCharOS

Antibodies-online

Antibody Resource

Biocompare

Labome

Any others?____________________________

8. Who decides which research antibodies to use in your research:

- Me
- Supervisor
- Post-doctoral colleagues in lab
- Other ____________________

9. Which of the following antibody-based methods have you used in your research:

- Western blotting
- Immunohistochemistry
- Immunohistochemistry – frozen sections
- Immunocytochemistry/Immunofluorescence
- Flow Cytometry
- Immunoprecipitation/ Co-immunoprecipitation
- Fluorescence-activated Cell Sorting
- Functional blocking assays
- ELISA

#### Appendix 7: Topic Guide for focus group

**Introduction**:

Welcome, introduction of researchers and project

Instructions regarding housekeeping:

- Please move to a quiet location without too much background noise and stable internet connection.
- If you disconnect from Teams during the focus group, please try to reconnect ASAP.
- If you have any technical difficulties during the focus group please use the chat function or email the researchers to alert us to any problems.
- The focus group will be 60-90 mins and will be recorded. Everything you say will remain confidential.
- After the session, the entire session will be transcribed word by word. The transcript will be checked and your details removed so the document is anonymous. Once this is completed, the original recording will be deleted.

Instructions regarding focus group:

- As you know, we have invited you to participate in this focus group because we are interested to know about how you choose and use antibodies in your research. We are also interested in your ideas for how to improve the process of choosing and using antibodies in research.
- We want to know about how you usually go about choosing an antibody for your experiments, and what you do next after you have chosen one.
- It's important to remember when answering and discussing questions that there are no right or wrong answers, please just be yourself and speak as honestly as possible. Also if you don't have an answer, then that's okay as well.
- Please be respectful of other participants' views and allow everybody the time to speak and voice their opinions.

Finally before we begin, are there any questions?

##### Main Questions

1. Please could you describe which area you work in, and what types of experiments you perform using antibodies?
2. We want to understand more about how researchers choose antibodies for their own research. Please could you describe how you go about choosing a new antibody?
   *Prompts:*
   - *Where do you search for antibodies?*
   - *What factors lead you to choose one antibody over another?*
3. In order to understand how information provided by antibody vendors influences decisions, we want to show you some information about 3 real antibodies raised against Annexin A11 (Appendix 8). Annexin A11 is thought to be expressed in the nucleus and cytoplasm of cells and has a predicted molecular weight of 54kDa. Variants in the gene encoding Annexin A11 are implicated in motor

neurone disease. These pictures and data are supplied by the antibody manufacturers. We will present the information on each antibody in turn.

a. Please discuss how you evaluate this information, whether you find it useful in deciding whether to buy the antibody.
b. How might you go about deciding which of these 3 to use? Which factors influence you?
c. For these antibodies, which applications do you think may or may not work, and why?
*Prompts:*
- *Does the data presented by the manufacturers support their use in immunofluorescence, western blotting or immunohistochemistry?*
- *Is there any additional data that you would have liked to see?*

d. Once you have decided on an antibody, what experiments (if any) would you perform to check it is working for your chosen application?
e. If the antibody did not work as you expected, what would be your next step?

4. What do you know about the five pillars of antibody validation?

5. The following figure explains the five main methods used to validate an antibody is performing adequately.
[Researcher goes through each in turn quickly summarising what they mean in practise]

Figure removed: "Antibody validation techniques"
(CiteAb; adapted from Uhlen et al., Nat Methods 2016).
Licence unverified - to be replaced with an original figure.

a. What barriers (if any) may prevent you or your lab from performing antibody validation experiments?
b. What do you think would help researchers choose and validate antibodies better in their research?

6. Do you have any other comments about antibodies and your personal experiences?

#### APPENDIX 8

### Antibody 10479-2-AP

| | |
|---|---|
| **Company** | **Proteintech** |
| **Cost** | **£269** |
| **Cost/100µL** | **£179** |
| **Host** | **Rabbit** |
| **Clonality** | **Polyclonal** |
| **Isotype** | **IgG** |
| **Clone number** | **N/A** |
| **Technique** | **WB, IHC, ICC/IF, IP/CoIP** |
| **Citations (CiteAb)** | **8** |

Manufacturer-supplied antibody image removed
(licence unverified)

Manufacturer-supplied antibody image removed
(licence unverified)

Manufacturer-supplied antibody image removed
(licence unverified)

Manufacturer-supplied antibody image removed
(licence unverified)

Manufacturer-supplied antibody image removed
(licence unverified)

Manufacturer-supplied antibody image removed
(licence unverified)

Manufacturer-supplied antibody image removed
(licence unverified)

Manufacturer-supplied antibody image removed
(licence unverified)

Manufacturer-supplied antibody image removed
(licence unverified)

Manufacturer-supplied antibody image removed
(licence unverified)

Manufacturer-supplied antibody image removed
(licence unverified)

Manufacturer-supplied antibody image removed
(licence unverified)

Manufacturer-supplied antibody image removed
(licence unverified)

Manufacturer-supplied antibody image removed
(licence unverified)

Manufacturer-supplied antibody image removed
(licence unverified)

Manufacturer-supplied antibody image removed
(licence unverified)

### Antibody ab236599

| | |
|---|---|
| **Company** | **Abcam** |
| **Cost** | **£350** |
| **Cost/100µL** | **£350** |
| **Host** | **Rabbit** |
| **Clonality** | **Polyclonal** |
| **Isotype** | **IgG** |
| **Clone number** | **N/A** |
| **Technique** | **WB, IHC, ICC/IF** |
| **Citations (CiteAb)** | **0** |

Manufacturer-supplied antibody image removed (licence unverified)

Manufacturer-supplied antibody image removed (licence unverified)

Manufacturer-supplied antibody image removed (licence unverified)

Manufacturer-supplied antibody image removed (licence unverified)

### Antibody PA5-96670

| | |
|---|---|
| **Company** | **Thermo Fisher Scientific** |
| **Cost** | **£366** |
| **Cost/100µL** | **£366** |
| **Host** | **Rabbit** |
| **Clonality** | **Polyclonal** |
| **Isotype** | **IgG** |
| **Clone number** | **N/A** |
| **Technique** | **WB, ICC/IF, IP/CoIP** |
| **Citations (CiteAb)** | **0** |

Manufacturer-supplied antibody image removed
(licence unverified)

Manufacturer-supplied antibody image removed
(licence unverified)

Manufacturer-supplied antibody image removed
(licence unverified)
