## Supplementary material for "Inadequate antibody validation places substantial numbers of animal and human tissue samples at risk of waste": S2 File

### Research antibody survey

The purpose of this study is to understand current antibody use in research, in order to find  ways to improve things for the research community. Your responses are entirely confidential and we do not collect personal information, other than your email, which will be stored separately from the survey for confidentiality. It would be most helpful to us if you could be as honest as possible in your responses.

**INFORMATION FOR PARTICIPANTS**

- This survey is only intended for people who use antibodies in their research.
- You will be asked to provide an institutional email address in order to participate.
- The data will be used by researchers at the University of Leicester to help improve the use of antibodies in research. The researchers are engaging with publishers, ethics boards, research funders and industry, including antibody vendors. Only anonymised data from this study will be shared with any of these. The researchers do not make or sell antibodies.
- Email addresses will only be used to confirm institutional association and to deliver Amazon vouchers to those who opt in to receive these for their participation (only 100 are available on first come basis)
- Email addresses will not be shared with any other individuals or organisations
- This research project will comply with the General Data Protection Regulations (GDPR, 2018)
- **Anonymised** data from this research project will be published and shared by the University of Leicester with the **anonymised** raw data deposited in our institutional Figshare repository.
- More information is available here:

https://drive.google.com/file/d/1NHdlmu4ak6IzvRlLOGkVAC-14FKg5srG/view?usp=sharing

* Required

1. Do you work for an institution or organisation (both public or private sector), that participates or contributes to biomedical research using antibodies? *

◯ Yes

◯ No

Consent Form

2. Do you consent to your data being processed as described in the INFORMATION FOR PARTICIPANTS above?

   Please close this browser window if you do not consent.
   *

   ◯ I consent to my data being processed

3. Please insert INSTITUTIONAL email below *

4. Do you wish to be considered to receive an Amazon voucher for your participation? The first 100 respondents will be eligible for a £5 amazon voucher which will be distributed by email.
   *

   ◯ Yes

   ◯ No

5. How long have you worked a lab environment? *

- ◯ <1 year
- ◯ 1 - 5 years
- ◯ 5 - 10 years
- ◯ >10 years

6. What statement best describes your research experience level? *

- ◯ Undergraduate level
- ◯ Masters level
- ◯ PhD student
- ◯ Post-Doc/Early career researcher
- ◯ Experienced Post-Doc
- ◯ Independent Investigator
- ◯ Other

7. Which of the following antibody-based methods have you used in your research? *

- [ ] Western blot
- [ ] Immunocytochemistry
- [ ] Immunohistochemistry
- [ ] ELISA
- [ ] Immunoprecipitation
- [ ] Flow cytometry
- [ ] Protein array
- [ ] Blocking/Neutralising assay
- [ ] Functional assay
- [ ] Proximity ligation assay
- [ ] Radioimmunoassay
- [ ] Gel shift
- [ ] Chromatin immunoprecipitation
- [ ] Reverse phase protein array
- [ ] Immunoelectron microscopy
- [ ] Other

8. Do you have experience selecting antibodies for research? *

- ○ Yes
- ○ No

9. Who decides the antibodies to use in your research? *

- [ ] Me
- [ ] Bench-supervisor
- [ ] Principal Investigator
- [ ] Other

10. Have you ever accessed or used any of the following antibody related resources? *

- [ ] CiteAb
- [ ] Antibodypedia
- [ ] The Human Protein Atlas
- [ ] YCharOS
- [ ] Antibodies-online
- [ ] Antibody Resource
- [ ] Biocompare
- [ ] Labome
- [ ] The ABCD database
- [ ] pAbmAbs
- [ ] Patented Antibody Database
- [ ] RRID Antibody Registry
- [ ] Biomed Resource Watch
- [ ] None of the above
- [ ] Other

11. For a new antibody raised against your protein of interest, how do you confirm selectivity? *

- [ ] Fluorescence minus one (FMO)
- [ ] Isotype control
- [ ] Secondary only control
- [ ] Blocking peptide
- [ ] CRISPR
- [ ] siRNA, shRNA
- [ ] Antibody independent assay e.g. Transcriptomics, proteomics
- [ ] Compare binding to an independent antibody
- [ ] tag the target protein (e.g. his-tag, GFP) and compare antibody staining between antibody of question and protein tag.
- [ ] Mass spectroscopy of the sample bound to the antibody
- [ ] Compare staining to a positive control cell line
- [ ] Compare staining to a negative control cell line
- [ ] None of the above
- [ ] Other

12. Are you aware of the "five pillars of antibody validation"? *

- ( ) Yes
- ( ) No

13. How important are these factors to you, on a scale of 1-5 (5 being most important), when choosing a new antibody for your research? *

| | 1 | 2 | 3 | 4 | 5 |
|---|---|---|---|---|---|
| Data in literature | ○ | ○ | ○ | ○ | ○ |
| Data within an antibody database | ○ | ○ | ○ | ○ | ○ |
| Citations in literature | ○ | ○ | ○ | ○ | ○ |
| Availability of a free sample | ○ | ○ | ○ | ○ | ○ |
| Antibody supplier data | ○ | ○ | ○ | ○ | ○ |
| Antibody reviews | ○ | ○ | ○ | ○ | ○ |
| Clonality | ○ | ○ | ○ | ○ | ○ |
| Previous use within your laboratory | ○ | ○ | ○ | ○ | ○ |
| Reputation of antibody supplier | ○ | ○ | ○ | ○ | ○ |
| Cost | ○ | ○ | ○ | ○ | ○ |

14. How important are these data sources on a scale of 1-5 (5 being most important) when selecting an antibody for any given application? *

| | 1 | 2 | 3 | 4 | 5 |
|---|---|---|---|---|---|
| Western blot - single band at the correct molecular weight | ○ | ○ | ○ | ○ | ○ |
| Knockout/ Knockdown controls | ○ | ○ | ○ | ○ | ○ |
| Immunohistochemistry staining pattern | ○ | ○ | ○ | ○ | ○ |
| Immunocytochemistry staining pattern | ○ | ○ | ○ | ○ | ○ |
| Tested using multiple cell lines | ○ | ○ | ○ | ○ | ○ |
| Highly cited within the literature | ○ | ○ | ○ | ○ | ○ |

15. On a scale of 1-5, how important do you think any of the below are in preventing scientists from being able to perform robust antibody quality control experiments (5 being most important)? *

| | 1 | 2 | 3 | 4 | 5 |
|---|---|---|---|---|---|
| Time | ○ | ○ | ○ | ○ | ○ |
| Money | ○ | ○ | ○ | ○ | ○ |
| Expertise | ○ | ○ | ○ | ○ | ○ |
| Lab equipment | ○ | ○ | ○ | ○ | ○ |
| Incentives | ○ | ○ | ○ | ○ | ○ |
| Awareness | ○ | ○ | ○ | ○ | ○ |
| Support from supervisor/PI | ○ | ○ | ○ | ○ | ○ |
| Lab culture | ○ | ○ | ○ | ○ | ○ |

16. How useful do you think the below might be in improving antibody reliability in research on a scale of 1-5 (5 being most useful)? *

| | 1 | 2 | 3 | 4 | 5 |
|---|---|---|---|---|---|
| Open data sharing | ○ | ○ | ○ | ○ | ○ |
| E-learning resources | ○ | ○ | ○ | ○ | ○ |
| Antibody workshops | ○ | ○ | ○ | ○ | ○ |
| Undergraduate lectures | ○ | ○ | ○ | ○ | ○ |
| Funding for antibody validation | ○ | ○ | ○ | ○ | ○ |
| Increasing the visibility of negative data | ○ | ○ | ○ | ○ | ○ |
| Publishers requiring antibody validation data | ○ | ○ | ○ | ○ | ○ |
| Third party validation services | ○ | ○ | ○ | ○ | ○ |
| Funders requiring antibody validation data | ○ | ○ | ○ | ○ | ○ |

17. If there are any further thoughts you would like to add about any of the above, please do so below:

Sorry you are not applicable for this survey

Microsoft Forms branding removed
